# SARS-CoV-2 modulates NK cell responses via induction of HLA-E and triggers expansion of adaptive NK cells during acute infection

**DOI:** 10.1101/2024.03.04.583260

**Authors:** Mohammad Zahidul Hasan, Maren Claus, Nadine Krüger, Sarah Reusing, Eline Gall, Christina Bade-Döding, Armin Braun, Carsten Watzl, Markus Uhrberg, Lutz Walter

## Abstract

HLA-E expression plays a central role for modulation of NK cell function by interaction with inhibitory NKG2A and stimulatory NKG2C receptors on canonical and adaptive NK cells, respectively. Here, we demonstrate that infection of human primary lung tissue with SARS-CoV-2 leads to increased HLA-E expression and show that processing of the peptide YLQPRTFLL from the spike protein is primarily responsible for the strong, dose-dependent increase of HLA-E. Targeting the peptide site within the spike protein revealed that a single point mutation was sufficient to abrogate the increase in HLA-E expression. Spike-mediated induction of HLA-E differentially affected NK cell function: whereas degranulation, IFN-γ production, and target cell cytotoxicity were enhanced in NKG2C^+^ adaptive NK cells, effector functions were inhibited in NKG2A^+^ canonical NK cells. Analysis of a cohort of COVID-19 patients in the acute phase of infection revealed that adaptive NK cells were induced irrespective of the HCMV status, challenging the paradigm that adaptive NK cells are only generated during HCMV infection. During the first week of hospitalization, patients exhibited a selective increase of early NKG2C^+^CD57^-^ adaptive NK cells whereas mature NKG2C^+^CD57^+^ cells remained unchanged. Further analysis of recovered patients suggested that the adaptive NK cell response is primarily driven by a wave of early adaptive NK cells during acute infection that wanes once the infection is cleared. Together, this study suggests that NK cell responses to SARS-CoV-2 infection are majorly influenced by the balance between canonical and adaptive NK cells via the HLA-E/NKG2A/C axis.

## INTRODUCTION

The severe acute respiratory syndrome coronavirus 2 (SARS-CoV-2) was first described at the end of December 2019 (*1*) and resulted in a pandemic of coronavirus disease 2019 (COVID-19). The efficient human-to-human transmission and the mutation rate of the virus led to the evolvement and spreading of several viral variants of concern (VOCs) that fueled the pandemic. The disease symptoms evoked by SARS-CoV-2 infection can vary substantially in patients, ranging from mild symptoms to severe disease with acute respiratory distress or multi-organ failure caused by pneumonia and hyperinflammatory immune responses associated with considerable mortality (*2*).

Natural Killer (NK) cells are known to mediate strong antiviral responses in the respiratory tract (*3*). These innate lymphocytes are activated by a balance of signals that are received from inhibitory and stimulatory receptors expressed on the surface of NK cells (*4*, *5*). Activation of NK cells result in release of cytokines and chemokines that stimulate and attract other immune cells as well as in direct killing of infected cells via degranulation of cytotoxic effector molecules such as granzyme B and perforin or release of TNF (*6–8*). However, SARS-CoV-2 virus employs several mechanisms to evade NK cell-but also T and B cell-mediated immune responses: several proteins of the virus are able to inhibit production of type I interferons in infected cells (*9*), mutations particularly of the receptor-binding region of the spike protein result in loss of binding of neutralizing antibodies (*10*, *11*). Furthermore, NKG2D ligands (*12*) as well as HLA class I proteins (*13–15*) are downregulated to evade NK cell and cytotoxic T cell responses.

An important and conserved family of NK cell receptors are the C-type lectin-like receptors NKG2A and NKG2C, which both form heterodimers with another C-Type lectin-like receptor, CD94. The NKG2A protein contains immunoreceptor tyrosine-based inhibitory motifs in the cytoplasmic tail and represents an inhibitory receptor, whereas NKG2C lacks such motifs and associates with the DAP12 adaptor molecule that harbors immunoreceptor tyrosine-based activation motifs and is a stimulatory receptor. Both CD94/NKG2A and CD94/NKG2C recognize target cells via HLA-E, but with different avidities: CD94/NKG2A has higher and CD94/NKG2C has lower avidity (*16*). HLA-E is a non-classical MHC class I protein that is associated with beta-2-microglobulin and presents peptides. The polymorphism of HLA-E is low and restricted to two major allotypes, HLA-E*01:01 and E*01:03, which differ in cell surface expression levels due to a dimorphism at amino acid position 107 (*17*).

A hallmark of HLA-E is its specificity for hydrophobic peptides (*18*). Important sources of such peptides are in healthy cells the leader sequences of classical HLA class I proteins and of the non-classical HLA-G (*19–21*). Shortage of appropriate peptide supply, such as for example seen in virus-mediated downregulation/destruction of HLA class I as part of their immune evasion strategy, results in inhibition of HLA-E release from the endoplasmic reticulum and decreased expression at the cell surface (*22*). In this ‘missing-self’ situation, canonical NK cells expressing CD94/NKG2A are activated due to a lack of inhibition by the downregulated HLA-E ligand (*19*, *23*).

Human cytomegalovirus (HCMV) has evolved a mechanism to avoid attack from canonical NKG2A^+^ NK cells by expressing a peptide in the UL40 protein that is TAP-independent and is able to bind and stabilize HLA-E, thereby restoring HLA-E cell surface expression (*24–26*). However, HLA-E is also recognized by NK cells expressing the stimulatory CD94/NKG2C receptor (*19*). As first described by the Lopez-Botet group, NKG2C^+^ NK cells are generated during acute HCMV infection and, in a subset of HCMV^+^ donors, remain present as long-living NKG2C^+^CD57^+^ clonal expansions during latency and possibly throughout life (*27–29*). NKG2C^+^CD57^+^ NK cells are generally characterized by expression of self-specific inhibitory KIR, highly efficient antibody-dependent cellular cytotoxicity (ADCC), and reduced responsiveness to cytokine stimulation due to downregulation of receptors for pro-inflammatory cytokines such as IL-12 and IL-18 (*29–34*). Due to their phenotypical adaptations, longevity, and ability to elicit specific recall responses, this kind of virus-specific memory NK cell is now commonly referred to as adaptive NK cell. Notably, similar cells were also identified in CMV-infected macaques (*35–37*). Moreover, a different type of long-living NK cell is also present in mice, recognizing MCMV-infected cells via recognition of the m157 protein by the stimulatory Ly49H receptor (*38*). Such convergent evolution in distantly related species such as primates and mice suggest a fundamentally important role for adaptive NK cells in control of infection and long-lasting latency.

It is currently unknown whether SARS-CoV-2 uses a similar strategy as HCMV to stabilize HLA-E expression in infected cells and whether adaptive NK cells are generated in the course of the infection process. One early study reported spike-mediated induction of HLA-E expression leading to inhibition of canonical NKG2A^+^ NK cells (*39*). More recently, an IFN-γ-mediated increase of HLA-E was observed in infected alveolar macrophages in a macaque model, again leading to inhibition of NKG2A^+^ NK cells (*40*). In contrast, one study reported that exogenous loading with a peptide from the NSP13 protein led to stimulation of canonical NKG2A^+^ NK cells (*41*). In the present study, we systematically investigated the role of SARS-CoV-2 for regulation of HLA-E expression on the level of primary lung tissue and of lung cell epithelial cells productively infected with virus, as well as by transfection of lung epithelial cells with candidate proteins from SARS-CoV-2 and analyzed their impact on NK cell function. We demonstrate that productive infection with SARS-CoV-2 leads to increased cell surface expression of HLA-E and identify the YLQPRTFLL peptide of the spike protein as immunodominant peptide mediating this effect. Upregulation of HLA-E expression by the SARS-CoV-2 spike protein on lung cells differentially modulated NK cell function with canonical NKG2A^+^ NK cells being inhibited and adaptive NKG2C^+^ being activated by the infected target cells. Analysis of hospitalized COVID-19 patients during the acute phase of infection revealed specific activation and expansion of early adaptive NK cells of the NKG2C^+^CD57^-^ phenotype whereas mature NKG2C^+^CD57^+^ adaptive NK cells remained unchanged. The present observations have important implication for the understanding of the role of the NK cell compartment in controlling SARS-CoV-2 infection.

## MATERIAL AND METHODS

### Human samples and ethics statement

Human peripheral blood from healthy volunteers was collected into EDTA-containing tubes from healthy volunteers after informed consent. Ethics statements for the blood collection were obtained from the Ethics Committees at the University Medical Center Göttingen (UMG) and Medical Faculty of Heinrich-Heine University Düsseldorf (HHU) accepted under application numbers 5/1/18 and 2019-383, respectively. PBMCs layer was separated from the peripheral blood using Pancoll human at a density of 1.077 g/ml (Pan Biotech) in a leucosep tube with gradient centrifugation (800 xg for 40 min). Carefully removed PBMCs layer was washed with cold DPBS and the isolated PBMCs pellet was stored at -140°C or overnight incubation in medium_PBMC1_ (RPMI-1640 + GlutaMax, 10% inactivated FBS, 0.1% Gentamicin, 500 U/ml IL-2 and 10 ng/ml IL-15) to carry out the experiment the next day.

Human lung lobes were acquired from patients who underwent lobe resection for cancer at KRH Klinikum Siloah-Oststadt-Heidehaus (Hannover, Germany) or Hannover Medical School (MHH, Hannover, Germany). These experiments were approved by the ethics committee of the Hannover Medical School (MHH, Hannover, Germany) and in compliance with *The Code of Ethics of the World Medical Association* (number 2701–2015). All patients or their next of kin gave written informed consent for the use of explanted lung tissue for research.

The study on acute and recovered COVID-19 patients was approved by the local ethics committees (#178, IfADo and #301-2008, CAPNETZ, Hannover Medical School). All participants gave informed consent. COVID-19 patients admitted to Klinikum Nord Dortmund between April 2020 and January 2021 were enrolled in the study. A total of 32 patients with samples from at least 3 visits (COV A-C) were included in the study. We further analyzed samples from 32 healthy controls (HC), which were collected in 2019 before the onset of SARS-CoV-2 pandemic, and 19 samples from subjects who had recovered (REC) from previous SARS-CoV-2 infection. REC samples were taken in March and April 2020 (55 ± 6.6 days after infection; range 48 - 71 days). Demographic information of all participants and clinical parameters of acute COVID-19 patients are shown in Claus et al. (*42*).

### Expression constructs used in this study

A plasmid for expression of the spike protein (Wuhan sequence) was kindly obtained from Markus Hoffmann (German Primate Center, Göttingen). The amino acid sequence is shown in **Supplementary figure S1**. Spike peptide YLQPRTFLL is localized at amino acid positions 269 to 277. In order to investigate the role of the YLQPRTFLL peptide in stabilizing expression of HLA-E, alanine substitutions were introduced at either position 270 or 273 to encode mutated spike constructs P2 (L270A) and P5 (R273A) (**Supplementary figure S1**). A further construct was established to express the spike protein with C-terminally tagged turboGFP (spike-turboGFP fusion protein), which is shown in **Supplementary figure S2**. We isolated mRNA from SARS-CoV-2-infected cells for reverse transcription in cDNA, which was used in PCR to amplify the coding region of the NSP13 gene. The NSP13 sequence was cloned in an expression vector (**Supplementary figure S3**).

### Transfection of cells and infection with SARS-CoV-2

Plasmid DNA was transfected into Calu-3 cells applying Lipofectamine 3000 following supplier’s instructions (Thermo Fisher Scientific). Plasmid DNA was also nucleofected into 721.221 cells using Lonza’s Kit-V by Nucleofector II electroporation-based transfection system according to the manufacturer’s instructions. Around 48-55 h post-transfection experiments were performed with the transfected cells. Calu-3 and A549 cells were infected with SARS-CoV-2, Pango lineage B.1.513 at an MOI of 0.01 for 1 h. After infection, the cells were washed with DPBS and further incubated in the respective cell culture medium in order to harvest the cell culture supernatant at 24 or 48 h post inoculation. Supernatant was applied to Vero E6 cells and plaques were counted to determine viral titers in terms of plaque-forming units per milliliter (pfu/ml) as previously described (*43*). Calu-3, 721.221, A549, T2 or Vero E6 cells were cultured in medium_Calu-3/A549_ (DMEM/F-12 + GlutaMAX+ Sodium Pyruvate + Sodium Bi-carbonate, 10% inactivated FBS and 0.1% Gentamicin), medium_721_ (RPMI-1640 + L-Glutamin + 25 mM HEPES, 10% inactivated FBS and 0.1% gentamicin), medium_T2_ (RPMI 1640, 10% inactivated fetal calf serum (FCS), 2mM L-Glutamine, 100 U/mL penicillin and 100 µg/mL streptomycin) or medium_VeroE6_ (DMEM supplemented with 5% FCS) and the incubation was maintained at 37°C with 5% CO_2_. All the plasmid DNA constructs encoding either spike protein, spike protein with mutation, turboGFP-tagged spike protein or NSP13 were transiently transfected into cells.

### Human precision-cut lung slices (PCLS) and infection with SARS-CoV-2

PCLS were generated as recently described (*44*). In brief, PCLS were maintained in medium_PCLS_ (DMEM/F12 without phenol red (Thermo Fisher Scientific), 1% penicillin-streptomycin, 50 mg/mL enrofloxacin and 250 µg/mL amphotericin) and vitality was determined by observing the movement of ciliated cells of the bronchi and bronchioles under light microscopy. PCLS with 60-100% cilia activity were then first infected with 1 × 10^5^ pfu of SARS-CoV-2 (Pango lineage B.1.513) viral particles for 1 h. Infected PCLS were washed and incubated for another 24 h in DMEM/F12 supplemented with 1% penicillin-streptomycin and mycotics. Supernatant was also harvested as described above and viral titers determined by titration on Vero-E6 cells prior to fixation with 4% PFA.

### Immunohistochemistry and confocal laser microscopy

PFA-fixed PCLS were processed through paraffin-embedded (FFPE) block and sectioned at 5 µm of thickness, mounted on the slide and immediately stored at 4°C before being deparaffinized in xylene and then rehydrated in descending alcohol concentrations following the program of Giotto automatic stainer (Diapath). These samples were boiled for 5 min in sodium citrate buffer (10 mM sodium citrate, 0.05% Tween 20 prepared in dH_2_0; pH 6). Mouse anti-human HLA-E (clone 4D12) primary antibody (MBL Lifescience) and then secondary antibody goat anti-mouse conjugated with APC (BioLegend) were applied with staining buffer (DPBS, 2% FCS and 2 mM EDTA) to detect cells expressing HLA-E. For detection of SARS-CoV-2-infected cells, PCLS sections were permeabilized with 1X intracellular staining permeabilization wash buffer (from 10X, BioLegend) to capture the nucleocapsid protein using a rabbit anti-SARS-CoV-2 nucleocapsid antibody (Sino Biological) followed by a goat anti-rabbit antibody conjugated with APC (Abcam). For the washing steps, PBST buffer (DPBS with 0.05% Tween 20) was used and a drop of Fluoromount-G mounting medium with DAPI (Thermo Fisher Scientific) was applied onto the PCLS sample for 5 min at RT before fixing the slide with a cover slip (24 Å∼ 24 mm). The Plan-Apochromat 63x/1.40 oil DIC M27 or plan-apochromat 10x/0.45 M27 objective in the confocal laser microscope LSM 800 (Carl Zeiss) was used to record the images. Analysis was performed with the ZEN 2.3 software (Carl Zeiss) for Windows and Fiji2 (ImageJ) for Mac OS was used to calculate the mean fluorescence intensity (MFI) selecting cells in a fixed area and then the MFI of the selected cells of the image was measured.

### Multicolor flow cytometry analysis

Calu-3 or A549 cells were harvested 24 or 48 h post-infection with SARS-CoV-2 using warm DPBS, and stained with mouse anti-human HLA-E primary antibody or mouse anti-human HLA class I primary antibody (clone W6/32, purified from hybridoma cells) in 1:100 dilution with cell staining buffer (BioLegend) and a secondary antibody goat anti-mouse antibody conjugated with APC or rat anti-mouse antibody conjugated with Alexa Fluro 700 (BioLegend). These stained cells were fixed with fixation buffer (BioLegend) prior to the flow cytometry experiment. Following same staining procedures but without fixation, Calu-3 or 721.221 cells were also stained for HLA-E expression 48-55 h post transfection of gene expression constructs.

NK cells were defined as positive for CD56^dim^ using Brilliant Violet 650 conjugated mouse anti-human CD56 (BioLegend) and PerCP/Cyanine5.5 conjugated mouse anti-human CD16 antibody (BioLegend) and negative for lineage CD3, CD14, CD19 and CD20 using mouse anti-human lineage cocktail 3 (lin 3, BD bioscience). NKG2A and NKG2C-expressing NK cells were identified using anti-NKG2A (clone Z199, Coulter) conjugated with APC and anti-NKG2C (clone 134591, R&D) conjugated with PE or AF700.

For degranulation assays mouse anti-human CD107a conjugated with PE (BD Biosciences) was added to the PBMCs incubated overnight in medium_PBMC1_ after washing with DPBS. These PBMCs were immediately added to the Calu-3 or 721.221 cells transfected with gene expression constructs or untransfected cells for 1 h at 37°C with 5% CO_2_. After 1 h of stimulation, 1X Monensin solution (from 1000X, BioLegend) and 1X Brefeldin A solution (from 1000X, BioLegend) was added and incubation continued for another 5 h prior to the staining of NKG2A or NKG2C expressing NK cells. First the percentage of CD107a+ cells within the NK cells was assessed, then also the percentage of CD107a+ NKG2A+ or CD107a+ NKG2C+ NK cells was further stratified. Similarly, the percentages of CD107a+ NK cells and NKG2A+ and NKG2C+ NK cell subsets were determined in PBMCs upon co-incubation with uninfected and SARS-CoV-2-infected Calu-3 cells.

For the detection of intracellular IFN-γ and TNF expression, PBMCs were stained for lineage (see above), CD56, NKG2A, and NKG2C before fixation and permeabilization with fixation buffer followed by cell permeabilization buffer (Triton X-100, Cell Signaling). These fixed and permeabilized cells were further stained with mouse anti-human IFN-γ antibody conjugated with PE (BioLegend) and Brilliant Violet 421 conjugated mouse anti-human TNF antibody (BioLegend). To exclude dead cells, staining with viability dye Zombie aqua or Zombie yellow (BioLegend) was also applied prior to flow cytometry experiments with spectral cell analyzer ID7000 (Sony). FlowJo version 10.8.0 in Mac OS operating system was used to analyze experimental data, to prepare the figures and to measure the MFI excluding the autofluorescence in each setup of experiment.

Sample preparation for multicolor spectral flow cytometry of acute and recovered patients’ samples is described in Claus et al. (*42*). Samples were analyzed on a 5 laser Cytek® Aurora (Cytek® Biosciences). Data were analyzed using the FlowJo software (version 10.8.2; FlowJo LLC, USA) incl. the plugins FlowAI, DownSample, tSNE (optSNE), Phenograph and Cluster Explorer for high parameter analysis.

### Western blot analysis

Calu-3 cells transiently transfected with SARS-CoV-2 spike (SP) or NSP13-encoding expression constructs or untransfected cells as negative control (neg. con.) were washed with ice cold DPBS and lysed in ice cold lysis buffer (50 mM Tris-HCl pH 7.5, 150 mM NaCl, 1% NP-40 (Fluka), 0.4 mM EDTA and 1 tablet protease-inhibitor-cocktail complete mini (Roche Diagnostic)) at 4°C with 25 min gentle shaking. Centrifugation (15 min, 16000 x g at 4°C) was performed to obtain the lysate. Lysate pre-cleaning steps were performed using Protein G Sepharose beads (GE Healthcare). To measure the protein content of the lysate a Qubit 4 fluorometer was used using the Qubit protein assay kit (Thermo Fisher Scientific). Equal amounts of pre-cleared lysate samples were separated in 8 – 16% Mini protean-TGX stain free protein gel (Bio-Rad) and the gel was transferred to a Trans-Blot Turbo Mini-PVDF membrane (Bio-Rad) for blotting using the Trans-Blot Turbo Transfer System (Bio-Rad). Everyblot Blocking Buffer (Bio-Rad) was used to block the membrane and to dilute the antibodies. For detection of HLA-E, 1:1200 diluted mouse anti-human HLA-E monoclonal primary antibody (clone MEM-E/02, Bio-Rad) and as secondary antibody goat anti-mouse polyclonal antibody conjugated with horse radish peroxidase (HRP) from Merck Millipore was used at a dilution of 1:3500. As loading control, HRP-conjugated rabbit anti-human GAPDH antibody (1:5000 dilution, Abcam) was used. Purified mouse anti-SARS-CoV-2 spike protein S2 antibody (1:3000 dilution, BioLegend) and HRP-conjugated goat anti-mouse secondary antibody (1:3750 dilution) or a purified rat anti-SARS-CoV-2 spike protein S1 antibody (1:1000 dilution, BioLegend) and HRP-conjugated goat anti-rat secondary antibody (1:3000 dilution, BioLegend) was applied for the detection of S1 or S2 subunits of SARS-CoV-2 spike protein in the transfected Calu-3 cells, respectively. NSP13 in transfected Calu-3 cells was detected using rabbit anti-SARS-CoV-2 NSP13 protein antibody (1:2000 dilution, St John’s Laboratory) with a goat anti-rabbit secondary antibody conjugated with HRP (1:8000 dilution, Santa Cruz Biotechnology). PageRuler Plus Prestained Protein Ladder (Thermo Fisher Scientific) was used to determine the molecular weight of protein samples separated on the gel. Finally, Immobilon Forte Western HRP Substrate (Merck Millipore) solution was applied to the blot to develop the signal for the chemiluminescent detection, and the image of the blot was acquired by ECL Chemocam Imager (Intas) equipped with ChemoStar imager software for Windows and the MFI of the targeted band was analysed using software Fiji2.

### Cell sorting and Annexin-V apoptosis assay

PBMCs were stained for lineage markers (see above) and CD56 (see above) in sorting buffer (PBS, 2% FCS and 2 mM EDTA) following the same staining protocol as described above. The CD56^dim^ NK cell population was sorted using a SH800 cell sorter (Sony) and collected immediately into medium_PBMC2_ (RPMI-1640 + GlutaMax, 10% inactivated FBS, 0.1% Gentamicin). These sorted cells were cultured for 1 h at 37°C with 5% CO_2_ with medium_PBMC2_ and then washed with DPBS prior to adding to the untransfected or transfected Calu-3 cells at 37°C with 5% CO_2_ for 8 h. After the incubation, cells were harvested and washed once with DPBS, and stained with Annexin-V-FITC with Propidium Iodide (PI) using Annexin-V kits according to the staining protocol of R&D system. The stained cells were immediately analyzed by flow cytometry and FlowJo was used to analyze the early and late apoptotic Calu-3 cells upon incubation with sorted NK cells. In addition, SARS-CoV-2 SP transfected Calu-3 cells were also incubated with mouse anti-human HLA-E antibody (4D12) for 1 h before adding the sorted NK cells for 6 h and subsequent analysis of Annexin V as described above.

### *In silico* analysis of HLA-E-binding peptides

*In silico* analysis of the complete proteome of SARS-CoV-2 (Wuhan-Hu1) was done with NetMHCpan 4.0 webserver at https://services.healthtech.dtu.dk/services/NetMHC-4.0/. Predictions were done for 9mer peptides of HLA-E alleles E*01:01 and E*01:03.

### Analysis of SARS-CoV-2 variants and mutation frequencies

The spike protein sequences of SARS-CoV-2 were analyzed for how often mutant sequences at single amino acid positions occur in the GISAID database available at https://gisaid.org/database-features/covsurver-mutations-app/usingthehCoV-19/Wuhan/WIV04/2019 as reference strain.

### Analysis of HCMV status

For qualitative determination of HCMV status in serum, a Cytomegalovirus (CMV) IgG ELISA (IBL International) was used according to the manufacturer’s instructions. CMV serostatus was determined relative to a cut-off control set to 10 units (U). Sample values < 9 U were defined as negative and values > 11 U were considered positive.

### Statistical analyses

All statistical analyses were conducted using the statistical analysis software GraphPad Prism 9 for Mac OS. The individual analyses are described in the respective figure legends.

## RESULTS

### Increased HLA-E expression on primary lung cells upon infection with SARS-CoV-2

It is well established that infection with SARS-CoV-2 results in downregulation of classical HLA class I proteins (*13–15*, *41*). However, whether HLA-E expression is also affected or changed at all is unclear. In order to investigate whether infection with SARS-CoV-2 results in changes of HLA-E cell surface expression, we analyzed human alveolar epithelium of human precision cut lung slices (PCLS). These lung slices were incubated with SARS-CoV-2 viral particles for 24 hours. Successful infection was verified by confocal microscopy showing the presence of SARS-CoV-2 nucleoprotein only in the infected lung slices (**figure 1A**). Further, infection with SARS-CoV-2 clearly enhanced cell surface expression of HLA-E (**figure 1B**, additional images shown in **Supplementary figure S4**). Quantitative analyses of single cells in the confocal microscopy images showed no differences for DAPI whereas HLA-E expression was significantly increased (p<0.0001) in infected PCLS compared to uninfected ones (**figure 1C**).

**Figure 1.**
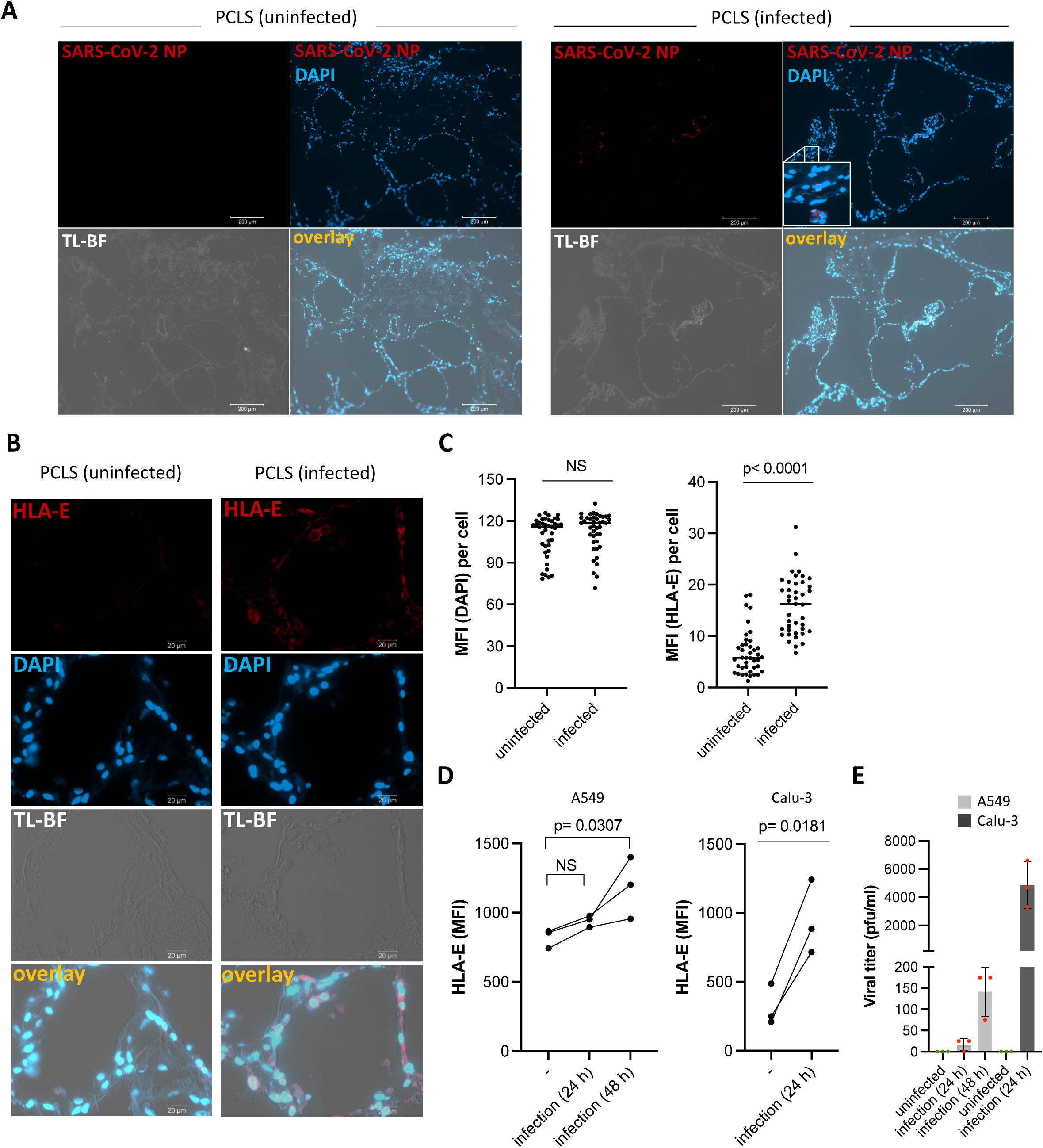
SARS-CoV-2 infection increases HLA-E expression. **(A)** Confocal laser scanning microscopy (CLSM) images of human precision-cut lung slices (PCLS) either uninfected (left panel) or infected with SARS-CoV-2, Pango lineage B.1.513, (right panel). PCLS were stained as indicated: presence of SARS-CoV-2 nucleocapsid protein (NP), cell nuclei (DAPI), and cell morphology (TL-BF) are shown in red, blue, and TL-BF channel recorded at plan-apochromat 10x/0.45 M27, respectively. Scale bar: 200 μm. **(B)** CLSM images of PCLS uninfected or SARS-CoV-2-infected stained with anti-HLA-E antibody (red) and stained for cell nuclei (DAPI, blue). Images were captured using plan-apochromat 63x/1.40 oil DIC M27 objective. Scale bar: 20 μm. **(C)** Single-cell mean fluorescence intensity (MFI) of DAPI (left panel) or HLA-E (right panel) was measured within the acquired images of uninfected or infected PCLS using ImageJ. Statistical analyses: t-test (parametric, unpaired, two-tailed, 95% confidence level), with p-values > 0.05 considered not statistically significant (NS). **(D)** Calculated MFI of HLA-E expression on uninfected or infected A549 cells at 24 h and 48 h post infection (left panel), and on uninfected or SARS-CoV-2 infected Calu-3 cells at 24 h post infection (right panel). Increasing the time post infection in Calu-3 resulted in a strong cytopathic effect and cell death due to high virus titers. Thus, only 24 h was measured as time point post infection. Statistical analyses: one-way ANOVA with Dunnett’s multiple comparisons test (uninfected A549 cells as control) or t-test (parametric, paired, two-tailed, 95% confidence level) for Calu-3 cells; p-values > 0.05 considered statistically NS. **(E)** Viral titers of SARS-CoV-2-infected or uninfected cells (A549 or Calu-3), quantified as plaque-forming units per milliliter (pfu/ml) applying the supernatant of uninfected or infected cells (Calu-3 or A549) on Vero E6 cells at 24 h or 48 h post infection. The graphs show mean and SD of three independent experiments.

The SARS-CoV-2-mediated induction of HLA-E expression could be further confirmed in two human lung adenocarcinoma epithelial cancer cell lines, Calu-3 and A549, which both serve as established cell line models to investigate infection of human lung tissue with coronaviruses (*45*). Infection of Calu-3 cells resulted in a significant increase of HLA-E on the cell surface after 24 h (**figure 1D**). In contrast, infection of A549 cells with SARS-CoV-2 showed no increase after 24 h but a slight, but statistically significant increase of HLA-E expression 48 h after infection (**figure 1D**). The kinetics of HLA-E induction in the infected cells (Calu-3 or A549) correlated well with SARS-CoV-2 virus titers in culture supernatant, which were much higher in Calu-3 compared to A549 cells (**figure 1E**). Due to the higher susceptibility to infection with SARS-CoV-2, Calu-3 cells were used for further experiments. Notably, in SARS-CoV-2-infected Calu-3 cells we could also confirm previous reports demonstrating decreased expression of classical HLA class I molecules (**Supplementary figure S5**). In summary, infection with SARS-CoV-2 led to upregulation of HLA-E cell surface expression on primary human PCLS as well as two different human lung cell lines.

### Induction of cell surface expression of HLA-E by the spike protein of SARS-CoV-2

Next, we aimed at unraveling the mechanisms for HLA-E upregulation and hypothesized that the effect could be mediated by a peptide encoded in the proteome of SARS-CoV-2 that would bind to HLA-E and stabilize its surface expression. As a starting point, we subjected the proteome of SARS-CoV-2 to *in silico* analysis to identify, in which proteins potential peptides might be located that bind to HLA-E with high affinity. Two proteins showed the highest prediction scores for HLA-E-binding peptides, the spike protein (YLQPRTFLL) and the NSP13 protein (VMPLSAPTL), confirming previous predictions by others (*39*, *41*, *46*) (**figure 2A**). In order to find out if these proteins could indeed upregulate HLA-E expression, we cloned full-length spike and NSP13 coding sequences in expression vectors for transfection in Calu-3 lung cells. Successful production and expected molecular weight of the spike protein with its S1 and S2 subunits as well as the NSP13 protein were confirmed by western blotting (**Supplementary figure S6**). Expression of the spike protein in Calu-3 cells led to increased HLA-E expression in flow cytometric analysis (**figure 2B**), accompanied by increased protein amounts in western blot analysis (**figure 2C**), which is generally expected for HLA class I when suitable peptides are available during quality control in the endoplasmic reticulum (*47*). Notably, untransfected Calu-3 cells already express HLA-E (genotype E*01:01, E*01:03, own unpublished data) due to expression of endogenous classical HLA class I allotypes (A*24, A*68; B*07, B*051; C*15), which supply HLA-E with fitting peptides. Therefore, we additionally included the 721.221 B-lymphoblastoid cell line in our experiments that is devoid of classical HLA class I expression (*48*) and consequently lacks endogenous HLA-E on the cell surface. Upon transfection of the spike protein-encoding construct in 721.221 cells, we detected significant induction of HLA-E on the surface of these cells, too (**figure 2D**). We also established a construct allowing for expression of a spike-turboGFP fusion protein to analyze if increasing amounts of spike protein correlate with higher expression of HLA-E. Indeed, increased HLA-E expression was found only in the GFP-positive cells and the amount of HLA-E on the cell surface strongly correlated with the amount of GFP, i.e. the amount of spike protein in each cell (**figure 2E**).

**Figure No 2.**
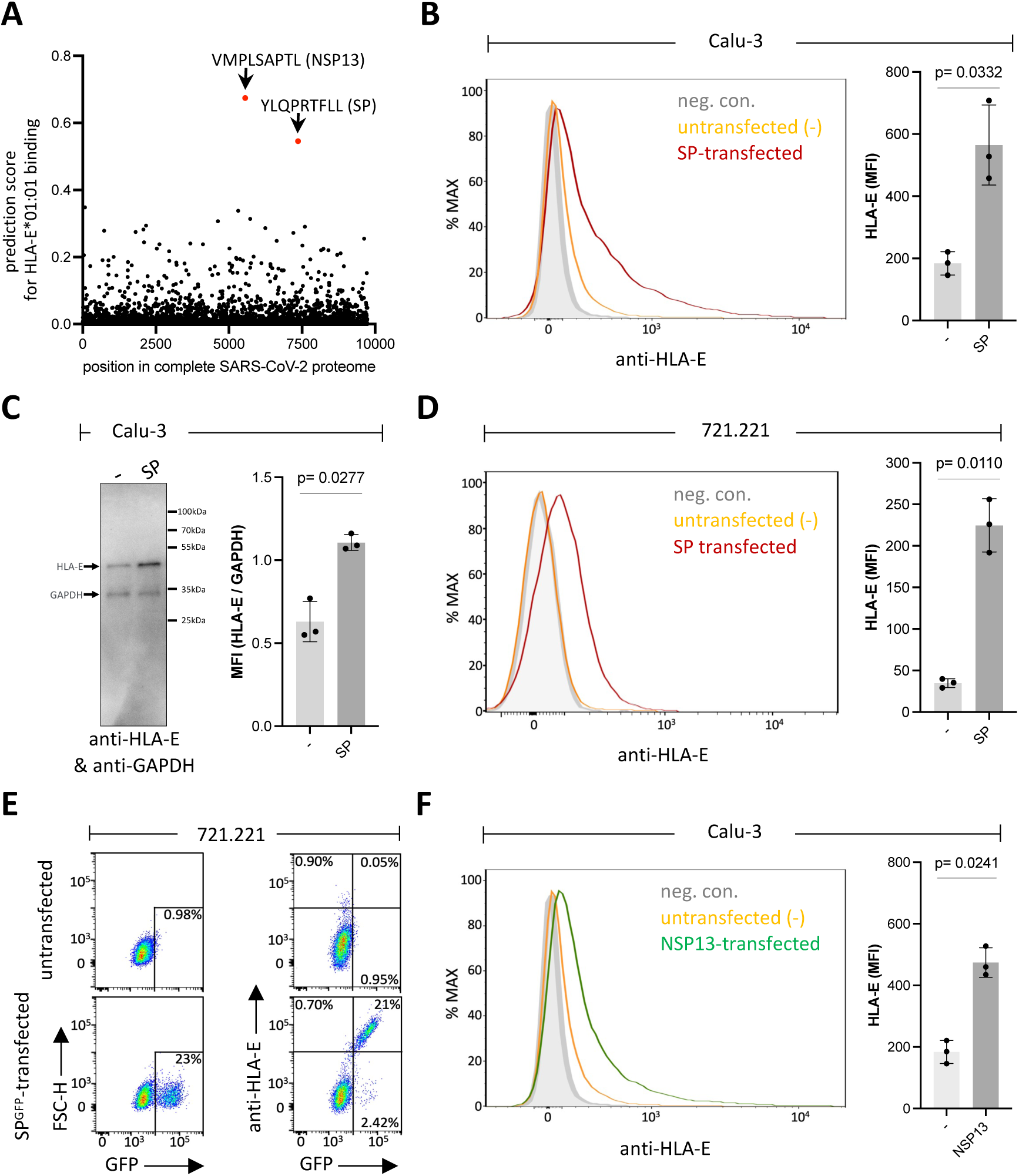
SARS-CoV-2 spike protein (SP) and non-structural protein 13 (NSP13) increase HLA-E expression. **(A)** *In silico* prediction of binding of SARS-CoV-2 protein-derived 9-mer peptides to HLA-E by analyzing the complete SARS-CoV-2 proteome (Wuhan-Hu1 genome sequence) using NetMHC-4.0. Peptides with a predicted high affinity for HLA-E are indicated. **(B)** Flow cytometric analysis of HLA-E expression at the surface of Calu-3 cells (left panel), either untransfected (yellow curve) or SP-transfected (red curve). Unstained (neg. con.) Calu-3 cells are shown as grey curve. Calculated MFI of HLA-E expression is displayed for untransfected (-) or SP-transfected Calu-3 cells (right panel). **(C)** Left panel shows western blot of HLA-E (∼45kDa) using anti-HLA-E antibody MEM-E/02 and cell lysates from untransfected (-) or SP-transfected Calu-3 cells. GAPDH was used as loading control (∼33kDa). Right panel shows normalized MFI of HLA-E obtained by dividing MFI of HLA-E by MFI of GAPDH of the same lysate. MFI was calculated on the detected HLA-E and GAPDH bands of untransfected (-) or SP-transfected Calu-3 cell lysates on WB analysis, n= 3 individual bands of separate experiments. **(D)** Left panel shows flow cytometric analysis of HLA-E expression at the surface of untransfected (-, yellow curve) or SP-transfected (red curve) 721.221 cells measured at 48-55 h after nucleofection. Unstained (neg. con.) 721.221 cells are shown as grey curve. The calculated MFI of cell surface HLA-E expression is indicated in the right panel. **(E)** Flow cytometry analysis of either only turboGFP (left panels) or turboGFP and HLA-E (right panels) in untransfected 721.221 cells or transfected with a construct for expression of turboGFP-tagged SP. Upper left quadrants in the right panels show HLA-E^+^GFP^-^ cells, while upper right quadrants show HLA-E^+^GFP^+^ cells, and lower right quadrants show HLA-E^-^GFP^+^ cells. Percentages are indicated. **(F)** The left panel shows histograms of HLA-E cell surface expression of untransfected (yellow curve) or NSP13-transfected (green curve) Calu-3 cells. Unstained (neg. con.) Calu-3 cells are shown as grey curve. The calculated MFI of cell surface HLA-E expression is indicated in the right panel. Statistical analyses: t-test (parametric, paired, two-tailed, 95% confidence level).

As outlined above, a second peptide with a high prediction score was found within the NSP13 protein. As shown in **figure 2F**, ectopic expression of the NSP13 protein led to an increase of HLA-E expression in Calu-3 cells that was comparable to that seen with the spike protein (**figure 2B**), thereby confirming that the peptide is efficiently processed from the NSP13 protein. Notably, in a previous study, the respective NSP13-derived peptide (VMPLSAPTL) was already shown to bind to HLA-E when exogenously loaded onto K562 cells. The peptide did however not lead to inhibition of NKG2A^+^ NK cells suggesting that it is not recognized by this inhibitory receptor (*41*) and might be regarded a nonfunctional peptide similar to what was described by Lin et al. (*49*). Here, we focused our further analyses on the spike protein and its effect on HLA-E and interaction with NKG2 NK cell receptors. In summary, we functionally confirmed the prediction from *in silico* analysis that the spike protein is able to stabilize HLA-E and thereby increases cell surface expression of this non-classical HLA class I protein.

### Spike protein stabilizes cell surface expression of HLA-E by providing the peptide YLQPRTFLL

To investigate the role of peptide YLQPRTFLL that is located at amino acid positions 269-276 and showed the highest predicted affinity in the spike protein, we used TAP-deficient T2 cells as established read-out system to investigate the effect of exogenously added peptides to stabilize cell surface expression of HLA-E. The T2 cell system was first tested for effectiveness. For this, we compared expression of the 721.221-AEH cells, which are known to strongly express HLA-E (*20*), with T2 cells expressing either the HLA-E*01:01 or the E*01:03 allotype and incubated with the HLA-G leader-derived peptide VMAPRTLFL (*50*), which can efficiently stabilize cell surface expression of HLA-E (*21*). The flow cytometric analysis showed the expected differences in HLA-E expression: AEH > E*01:03^VMAPRTLFL^ > E*01:01^VMAPRTLFL^ > control (**figure 3A**). Notably, the well-known expression differences of the two HLA-E allotypes to be observed on human primary cells were also reflected here in this T2 cell read-out system. Additionally, HLA-E cell surface expression correlated well with the amount of added VMAPRTLFL positive control peptide (**figure 3B**). Using the T2 system, we investigated the respective spike peptide YLQPRTFLL in comparison to peptides that were in silico predicted to bind to HLA-E with low affinity (INITRFQTL; position in spike 233-241), or an irrelevant peptide (YEQFAYDGK; position 113-121 in HLA-E) predicted to not bind to HLA-E, as well as the HLA-G derived peptide from the HLA-G leader sequence (VMAPRTLFL), serving as positive control. Spike peptide YLQPRTFLL and the positive control revealed comparably strong upregulation of HLA-E cell surface expression in T2 cells transfected with HLA-E*01:03, while the low affinity peptide INITRFQTL as well as the irrelevant peptide failed to induce HLA-E (**figure 3C**). T2 cells transfected with the HLA-E*01:01 allotype did only weakly discriminate between the different peptides, which is probably due to the generally lower level of HLA-E induction compared to the HLA-E*01:03 transfectant (**figure 3C**). In summary, spike peptide YLQPRTFLL has a strong stabilizing effect on cell surface expression of HLA-E, in particular of the E*01:03 allotype.

**Figure 3.**
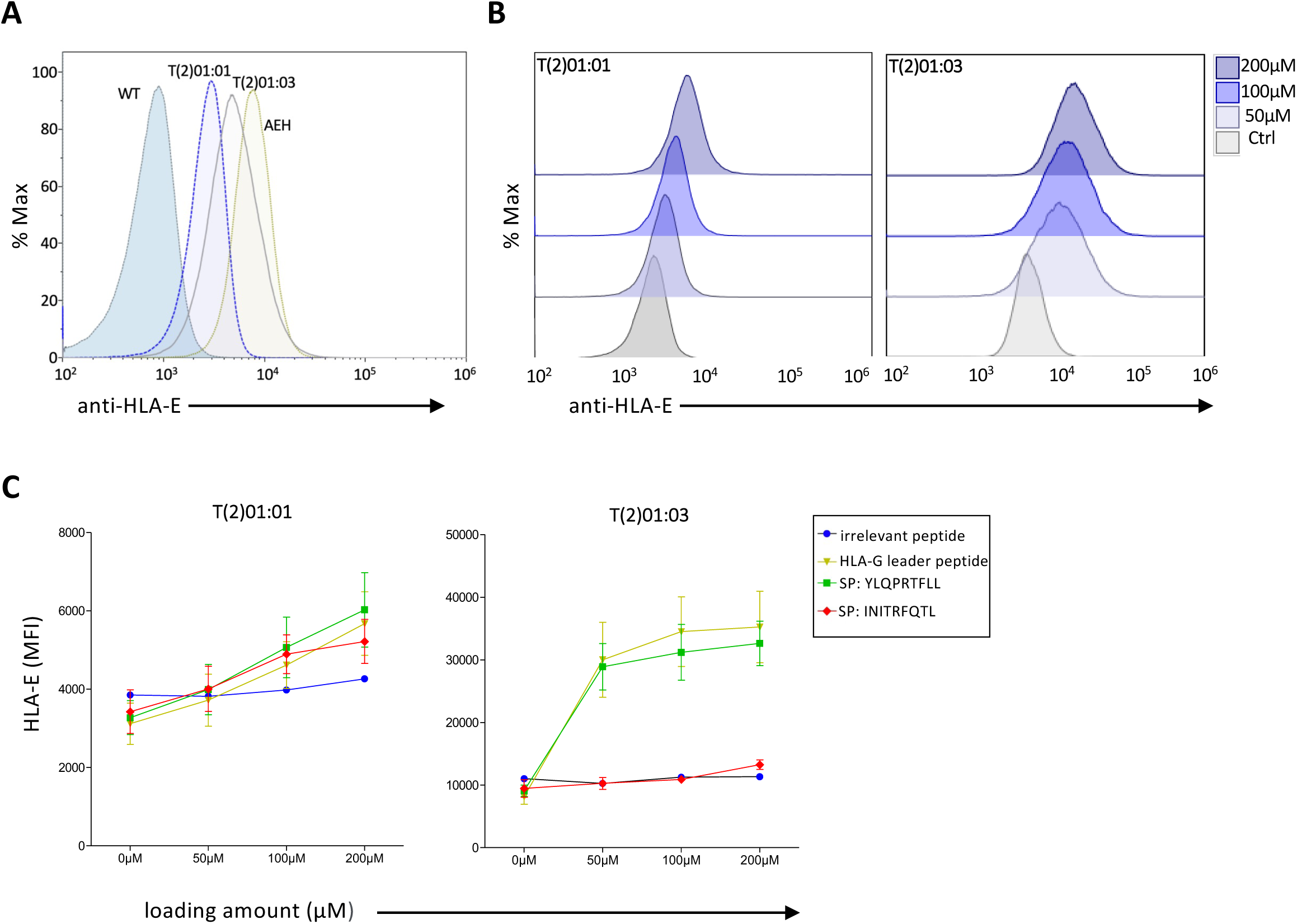
SARS-CoV-2 spike protein-derived YLQPRTFLL peptide stabilizes HLA-E cell surface expression. **(A)** Flow cytometric analysis of HLA-E cell surface expression on TAP-deficient T2 cells expressing either HLA-E^∗^01:01 or HLA-E^∗^01:03 and loaded with 200 μM of HLA-G leader peptide VMAPRTLFL in comparison to untransfected T2 cells without peptide loading (WT) and 721.AEH cell line strongly expressing HLA-E (*20*). **(B)** Flow cytometry data of HLA-E^∗^01:01 (left panel) or HLA-E^∗^01:03 (right panel) expression on T2 cells loaded with different concentrations (0−200 μM) of HLA-G peptide. **(C)** MFI of HLA-E^∗^01:01 (left panel) or HLA-E^∗^01:03 (right panel) on T2 cells loaded with the indicated peptides from SARS-CoV-2 spike, HLA-G leader peptide, or irrelevant peptide.

### Site-specific mutation of the spike protein abrogates induction of HLA-E expression

Although we could show that exogenously added YLQPRTFLL efficiently stabilizes expression of HLA-E, its contribution in comparison to other naturally processed peptides from the spike protein was so far unknown. To this end, we mutated the spike protein at single amino acid positions by introducing alanine residues at either position P2 (…Y**A**QPRTFLL…) or P5 (…YLQP**A**TFLL…) of the (**figure 4A**), which are known to be important residues for binding of peptides to HLA-E (*51*). Expression of the wildtype and the two mutant spike constructs in Calu-3 lung cells clearly indicated that single amino acid changes in the spike protein significantly changed HLA-E expression resulting in the HLA-E expression order: spike wildtype > spike P2 mutant > spike P5 mutant (**figure 4B**). In fact, mutation of P5 completely abrogated the spike-mediated increase of HLA-E expression, strongly suggesting that increased surface expression of HLA-E is largely dependent on processing and loading of the YLQPRTFLL peptide (**figure 4B**). Mutation at the P2 position of YLQPRTFLL had an intermediate effect, but resulted in still higher expression of HLA-E as compared to untransfected Calu-3 cells (p<0.05; **figure 4B**). Together the data largely exclude the presence of additional major HLA-E-stabilizing peptides within the spike protein.

**Figure 4.**
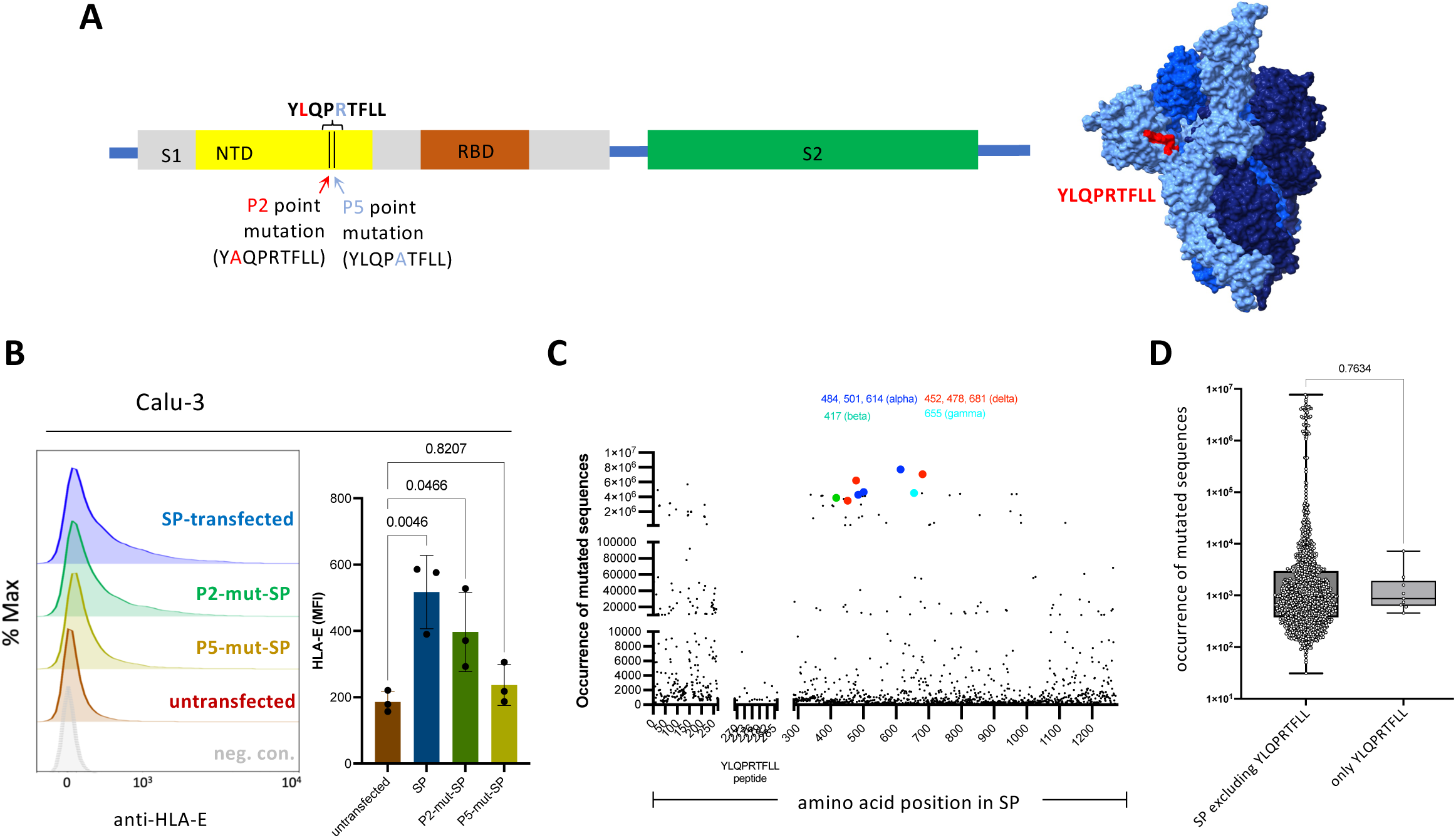
SARS-CoV-2 spike peptide YLQPRTFLL is responsible for increased HLA-E cell surface expression. **(A)** Scheme of spike (SP) protein gene expression construct with subunits S1 shown in grey and S2 shown in green. N-terminal domain (NTD) and receptor binding domain (RBD) are indicated in S1. Point mutations at position 2 (P2) and at position 5 (P5) of YLQPRTFLL are illustrated with corresponding mutated sequences. A three-dimensional model of the spike protein trimer is shown with location of the YLQPRTFLL peptide. **(B)** Left panel shows histograms of HLA-E expression at the surface of Calu-3 cells transfected with SP wildtype (violet), P2-mutant-SP (L270A, green), P5-mutant-SP (R273A, yellow), untransfected (red), and unstained cells (grey), respectively. The corresponding MFI is displayed in the right panel. Statistical analyses: one-way ANOVA with Dunnett’s multiple comparisons test (using untransfected Calu-3 cells as control). **(C)** Frequencies of mutated amino acid positions in spike protein sequences of the GISAID SARS-CoV-2 variant database as compared to the firstly described Wuhan-Hu1 SARS-CoV-2 spike sequence are shown. Key mutant residues characterizing the known SARS-CoV-2 variants of concern (alpha, beta, delta, gamma) are indicated. **(D)** Frequencies of mutated amino acid positions in the spike protein with exclusion of the YLQPRTFLL peptide and in the YLQPRTFLL peptide only are shown. The mean frequencies are not different. Statistical analysis: t-test (parametric, unpaired, two-tailed, 95% confidence level).

In order to find out whether the region encoding spike peptide YLQPRTFLL in the SARS-CoV-2 genome is subject to frequent mutation, we investigated a public database (GISAID) on frequencies of mutated SARS-CoV-2 amino acid positions. We found that the respective spike peptide is clearly not a target of frequent mutation (**figure 4C**) and is comparable to the mean mutation frequency within the spike protein (**figure 4D**). Importantly, the sequence remains conserved in all common SARS-CoV-2 variants of concern that evolved over the last four years (see CoVsurver mutations app at gisaid.org), confirming similar findings published previously by others (*46*). This sharply contrasts with other parts of the spike protein, where the common SARS-CoV-2 variants show extremely high mutation frequencies (**figure 4C**).

### Spike protein-mediated induction of HLA-E differentially affects NK cell function

NK cells are the main lymphocyte population interacting with HLA-E via their specific receptors: on the one hand canonical NK cells expressing the inhibitory CD94/NKG2A receptor and being the largest NK cell population in HCMV^-^ donors, and on the other hand adaptive NK cells, present in HCMV^+^ donors and expressing high levels of the stimulatory CD94/NKG2C receptor. In order to unravel how modulation of HLA-E expression by the spike protein of SARS-CoV-2 affects specific NK cell responses against infected target cells, we co-incubated Calu-3 cells that were either left untransfected or transiently transfected with spike protein and measured mobilization of cytotoxic granules to the cell surface (CD107a) as well as expression of the NK cell effector cytokines IFN-γ and TNF. In HCMV^-^ donors, spike protein-induced upregulation of HLA-E on the Calu-3 target cells led to significant reduction of degranulation (**figure 5A**, **upper panels; gating strategy is shown in Supplementary figure S7**) as well as IFN-γ expression (**figure 5B**, **upper panels**) in NK cells, indicating broad functional inhibition of canonical NK cells. Analyses of NKG2A^+^ and NKG2C^+^ subpopulations revealed that the large subpopulation of NKG2A^+^ NK cells was strongly inhibited while no differences in NK cell function were noticed in the small subpopulation of NKG2C^+^ NK cells upon incubation with untransfected and transfected target cells (**figure 5A**, **upper panels**). Of note, NKG2C^+^ NK cells in HCMV^-^ donors do express only low levels of NKG2C and these are known to be less functional than adaptive NK cells of HCMV^+^ donors, likely explaining their unresponsiveness to spike-induced upregulation of HLA-E expression in our experimental setup. Finally, the expression of TNF did not change in either NK cell population in HCMV^-^ donors (**figure 5C**, **upper panels**).

**Figure 5.**
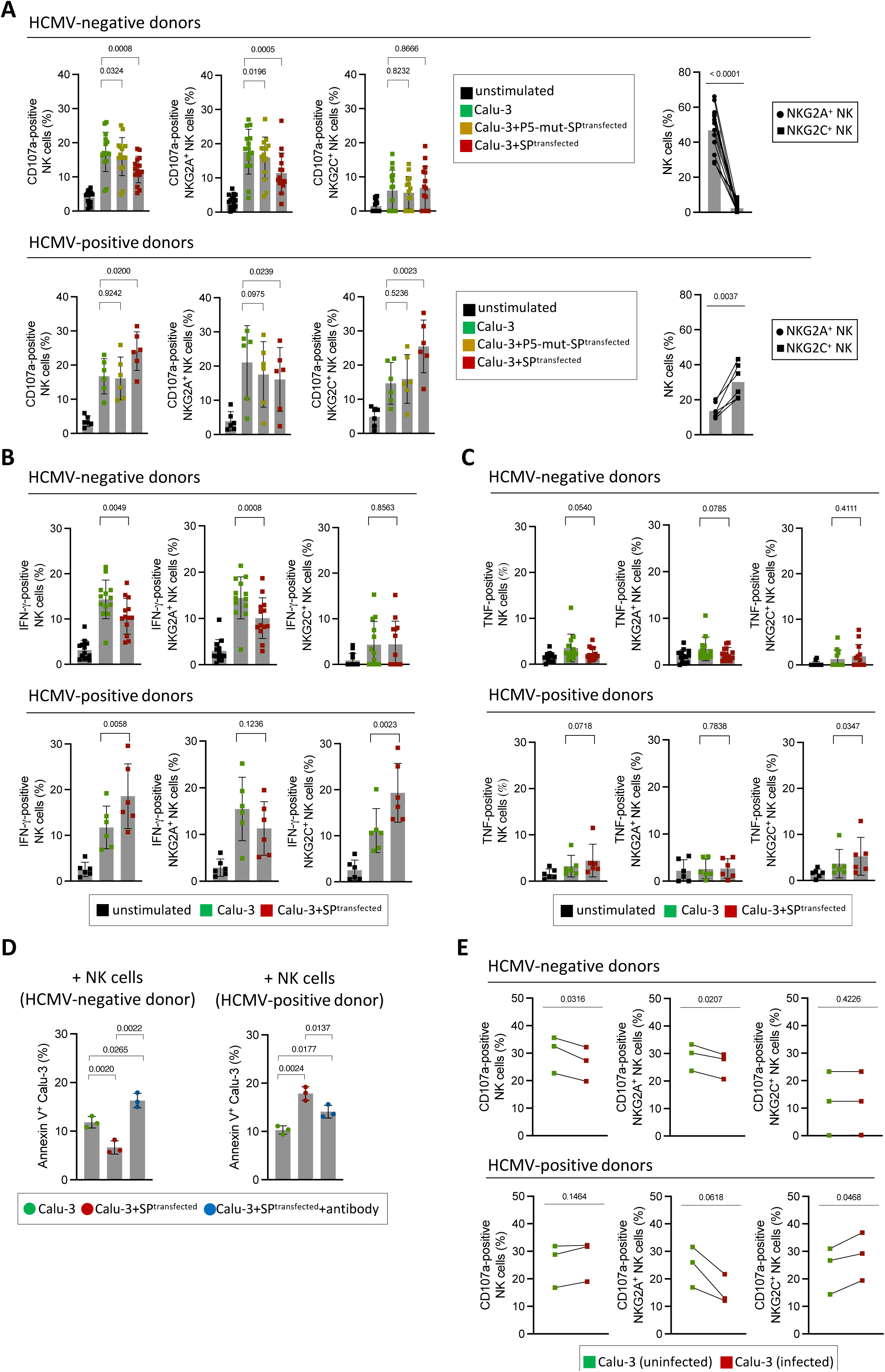
NK cell responses to cells expressing SARS-CoV-2 spike. **(A)** Degranulation of NK cells and NKG2A^+^ and NKG2C^+^ subsets from of PBMCs of HCMV^-^ donors (upper panels) and HCMV^+^ donors (lower panels). CD107a was measured in untreated NK cells or upon co-incubation with Calu-3 cells either untransfected or transfected with expression constructs encoding spike (SP) or spike with single point mutation at position R273A (P5-mutant SP). For this assay, n=15 HCMV^-^ donors or n= 6 HCMV^+^ donors were utilized. The right panels show frequencies of NKG2A+ and NKG2C+ NK cells in HCMV^-^ (right upper panel) and HCMV^+^ (right lower panel) donors. Statistical analyses: one-way ANOVA with Dunnett’s multiple comparisons test; using untransfected Calu-3 as control; student’s t-test (parametric, unpaired, two-tailed, 95% confidence level). Statistical analyses: one-way ANOVA with Dunnett’s multiple comparisons test; using untransfected Calu-3 as control; t-test (parametric, unpaired, two-tailed, 95% confidence level). Gating strategy for multicolor flow cytometric analysis is available in **Supplementary figure S7. (B)** Percentages of IFN-γ or **(C)** TNF-positive NK cells and NKG2A^+^ and NKG2C^+^ NK cell subsets were determined upon co-incubating PBMCs (n=13 HCMV^-^ and n=6 HCMV^+^ donors) with untransfected or SP-transfected Calu-3 cells as indicated. Percentages of IFN-γ or TNF-positive untreated NK cells shown as unstimulated control. Statistical analyses: t-test (parametric, paired, two-tailed, 95% confidence level). **(D)** Sorted NK cells from randomly chosen donors were co-incubated for 8 h with either untransfected or spike (SP)-transfected Calu-3 cells at an effector:target ratio of 2:1. Percentages of Annexin-V^+^, i.e. early apoptotic, cells are indicated. Early apoptosis on SP-transfected Calu-3 cells was also measured after blocking of HLA-E with an anti-HLA-E antibody (clone 4D12). Gating strategy for multicolor flow cytometric analysis as well as early and late apoptotic cells are shown in **Supplementary figure S8**. Statistical analysis: one-way ANOVA with Tukey’s multiple comparisons test comparing the mean of each condition. **(E)** Percentages of CD107a+ NK cells as well as NKG2A+ and NKG2C+ NK cell subsets from n=3 HCMV^-^ donors and from n= 3 HCMV^+^ donors upon co-incubation with SARS-CoV-2-infected Calu-3 cells measured 24 h after infection. Statistical analyses: t-test (parametric, paired, two-tailed, 95% confidence level). Assays shown in (A-C, E) were performed at a PBMC:target cell ratio of 5:1.

Analysis of NK cells in HCMV^+^ donors, which had variably sized expansions of adaptive NKG2C^+^ NK cells (**figure 5A**, **lower panels**), exhibited a very different functional outcome. Spike-mediated HLA-E upregulation on Calu-3 cells led to increased degranulation (**figure 5A**, **lower panels**) and IFN-γ (**figure 5B**, **lower panels**) production of NK cells. Analyses of NKG2A^+^ and NKG2C^+^ subpopulations in these donors revealed a strong increase in degranulation in the large subpopulation of NKG2C^+^ adaptive NK cells, while the smaller subpopulation of NKG2A^+^ NK cells showed less degranulation (**figure 5A**, **lower panels**) and a non-significant decrease in IFN-γ production (**figure 5B**, **lower panels**) upon co-incubation with spike-expressing Calu-3. The activation of NKG2C^+^ adaptive NK cells was also mirrored in a slight but statistically significant increase of TNF expression in these donors, which was also seen as a trend (p=0.0718) in overall NK cells (**figure 5C**, **lower panels**). The immunodominant role of the YLQPRTFLL sequence within the spike protein could be confirmed by comparative analysis of spike and mutant spike protein (P5 mutant, see figure 4A): the stimulatory effect of the spike protein on degranulation of adaptive NK cells in HCMV^+^ donors was completely abolished (**figure 5A**, **lower panels**) and the inhibitory effect on canonical NK cells in HCMV^-^ donors majorly reduced (**figure 5A**, **upper panels**).

Besides investigating degranulation in NK cells, we also examined induction of apoptosis in spike protein transfected target cells using annexin V and propidium iodide staining (**figure 5D**, **Supplementary figure S8**). When using flow cytometrically sorted NK cells from an HCMV^-^ donor, early apoptosis was significantly decreased in spike-transfected Calu-3 target cells compared to untransfected ones, reflecting NK cell inhibition of canonical NK cells through the NKG2A receptor (**figure 5D**, **left panel).** In contrast, when sorted NK cells from an HCMV^+^ donor were taken, apoptosis of spike-transfected target cells was increased, reflecting stimulation of adaptive NK cell effector function by increased HLA-E expression (**figure 5D**, **right panel**). Finally, blocking of HLA-E/peptide accessibility using an HLA-E antibody led to a strong increase of target cell apoptosis in case of NK cells from the HCMV^-^ donor and vice versa inhibition of adaptive NK cells from the HCMV^+^ donor (**figure 5D**). As expected, in the latter case the decrease in apoptosis was only partial, reflecting the variability in composition of the NK cell compartment in HCMV^+^ donors that is a mixture of adaptive NK cells that get stimulated and canonical NK cells that get inhibited. Further, the antibody blocking experiments provided formal proof of HLA-E being the receptor ligand responsible for the observed functional outcome on NK cells.

Recognition by NKG2A^+^ and NKG2C^+^ NK cells is not only dependent on the strength of HLA-E expression, but also on competition between functional and nonfunctional peptides (*49*). Hence, in productively infected cells, the functional effects elicited by the HLA-E/YLQPRTFLL complex could possibly be neutralized by competing nonfunctional peptides such as the recently described peptide from the NSP13 protein (*41*). To find out if this is the case, Calu-3 cells were successfully infected with SARS-CoV-2 and 24 h post infection the infected cells were co-cultured with PBMCs to assess NK cells’ function. As expected, in HCMV^-^ donors degranulation of NKG2A^+^ NK cells was reduced, whereas no change was noticed for NKG2C^+^ NK cells, leading to overall inhibition of the NK cell compartment (**figure 5E**, **upper panels**). In contrast, in HCMV^+^ donors, degranulation of NKG2C^+^ adaptive NK cells was significantly increased, while degranulation of NKG2A^+^ NK cells was decreased (**figure 5E**, **lower panels**). Together, the functional consequences of productive SARS-CoV-2 infection were comparable to those elicited by the spike protein alone, suggesting that the spike protein, or rather the spike-encoded peptide YLQPRTFLL, is indeed dominantly modifying NK cell function in SARS-CoV-2 infection.

In summary, we demonstrate that SARS-CoV-2-induced HLA-E expression on infected lung cells has a differential impact on virus-specific NK cell responses arising from the balance between inhibition of canonical NKG2A^+^ cells and stimulation of adaptive NKG2C^+^ NK cells.

### Activation of adaptive NK cells in acute SARS-Cov-2 infection is independent of HCMV status

The above observations suggested a substantial influence of adaptive NK cells on the overall NK cell response against SARS-CoV-2 infected cells. Given the similarities to HCMV infection with regard to the opposing modulation of classical HLA class I and HLA-E expression, we wondered whether SARS-CoV-2 infection would trigger *de novo* generation of adaptive NKG2C^+^ NK cells. To this end, we analyzed a cohort of COVID-19 patients that were hospitalized between April 2020 and January 2021 and that were already shown to exhibit an infection-related increase of NKG2C^+^ adaptive NK cells (*42*). In order to distinguish between newly arising and already established adaptive NK cells, we analyzed the presence of CD57, a carbohydrate epitope that accumulates during terminal differentiation of NK cells. As outlined in **figure 6A**, the frequency of mature adaptive NK cells (NKG2C^+^CD57^+^) did not change in the acute phase of SARS-CoV-2 infection, neither in HCMV^+^ nor in HCMV^-^ patients. In contrast, the frequency of early adaptive NK cells (NKG2C^+^CD57^-^) increased in the majority of HCMV^+^ and HCMV^-^patients (**figure 6B)**. Analysis of HCMV-specific IgM titers (data not shown) excluded that any of the HCMV^-^ donors was experiencing a primary HCMV infection at an early phase before IgG seroconversion is detectable. In one HCMV^-^ case (patient #49), the frequency of early adaptive NK cells increased more than 5-fold in the 7d observation period since admission to the hospital, whereas the frequency of mature adaptive NK cells remained low **(figure 6C)**. There was also one HCMV^-^ case (patient #18) that showed a decrease in frequency of early adaptive NK cells and this was accompanied by a strong decrease of the activation marker HLA-DR, whereas the other five HCMV^-^ patients showed a strong increase **(Figure 6D)**, making it likely that patient #18 was already in the contraction phase of the adaptive NK cell response. This is also consistent with the fact, that this patient had the highest initial frequency of early adaptive NK cells (3.6%) compared to the other HCMV^-^ patients (1.6% ± 1.3%).

**Figure 6.**
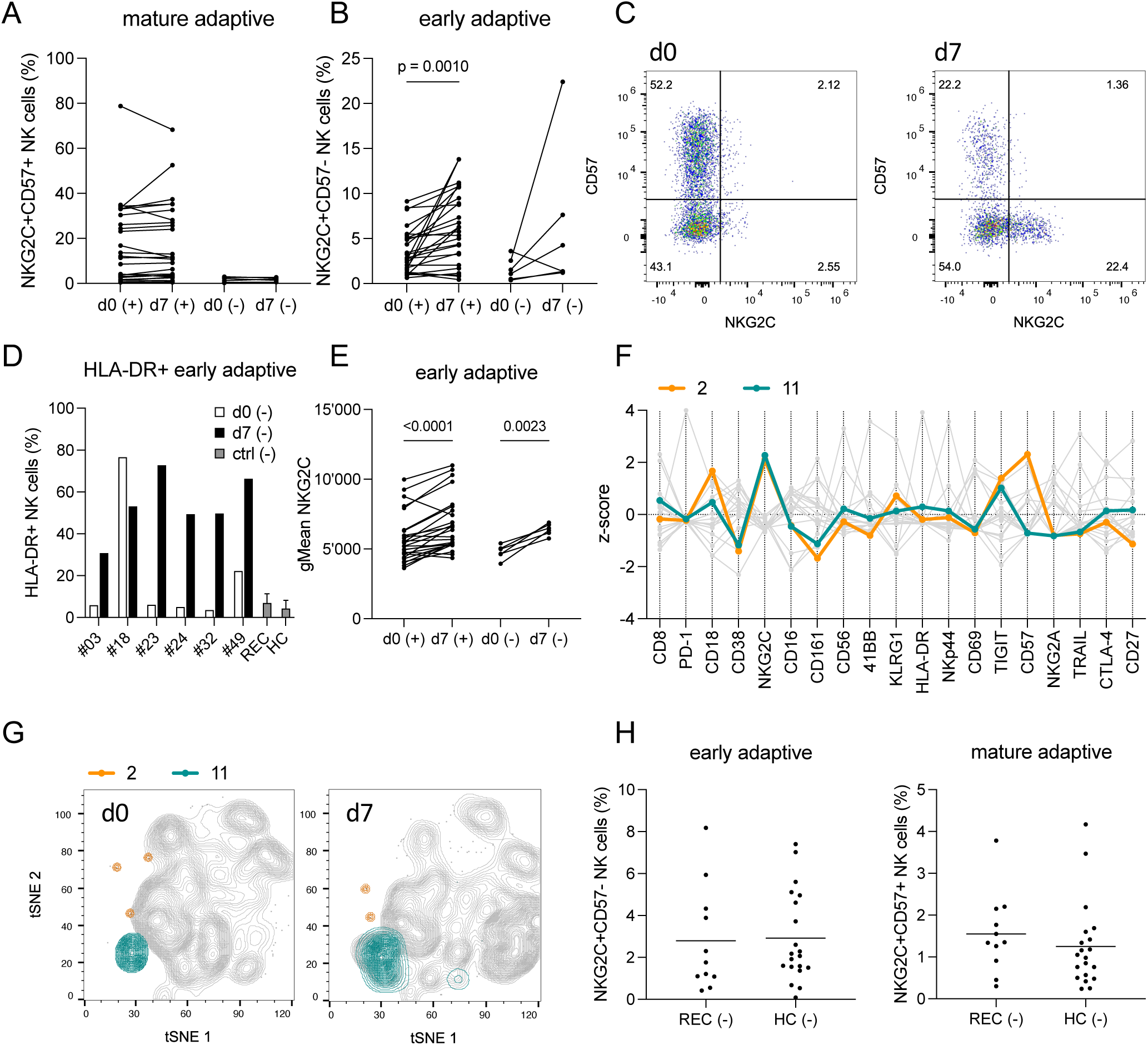
Emergence of early adaptive NK cells in acute SARS-Cov-2 infection is independent of HCMV status. Percentages of **(A)** mature adaptive NKG2C+CD57+ and **(B)** early adaptive NKG2C+CD57-NK cells from HCMV (+) and (-) patients at day 0 and day 7. **(C)** Representative data of one individual HCMV (-) patient (#49) showing significant expansion of early adaptive NK cells from timepoint d0 to d7. **(D)** Percentage of HLA-DR+ early adaptive NK cells from HCMV (-) acute Covid-19 patients at timepoints d0 and d7, recovered patients (REC) and healthy controls (HC). **(E)** Cell surface expression level (geometric mean) of NKG2C on early adaptive NK cells from HCMV (+) and (-) patients at timepoints d0 and d7. **(F)** Features of Phenograph clusters #2 and #11 representing NKG2C+CD57+ (#2) and NKG2C+CD57-(#11) NK cells. Data are shown as z-score of MFI. **(G)** tSNE plots of one representative HCMV (-) patient highlighting clusters #2 and #11 at timepoints d0 and d7. **(H)** Percentage of early adaptive and mature adaptive NK cells from HCMV (-) healthy controls (HC) and recovered patients (REC). Statistical analyses: Kruskal-Wallis test with Dunn’s multiple comparisons test and Mixed-effects analysis with Šídák’s multiple comparisons test. Legends of supplementary figures are included in the respective figures S1 to S10.

All HCMV^-^ and most HCMV^+^ patients exhibited a pronounced increase of NKG2C expression levels on early adaptive NK cells from day 0 to day 7 **(figure 6E)**, further illustrating their consistent activation. The increase of early adaptive NK cells during acute infection was further illustrated by Phenograph analysis of the high-dimensional flow cytometry data, which identified 2 distinct clusters resembling mature adaptive (cluster #2) and early adaptive (cluster #11) NK cells, respectively **(figure 6F)**. While the frequency of cluster #2 did not change during the course of infection, cluster #11 increased in both HCMV+ and HCMV-patients from day 0 to day 7 **(figure 6G**, **Supplementary Figure S9)**.

Finally, since we had no long-term follow up of the acute COVID-19 patient cohort, we utilized an independent cohort of recovered (REC) patients (55d ± 6.6d after infection) to address the question whether the observed expansion of early adaptive NK cells eventually leads to mature adaptive NK cells that remain present also in HCMV^-^ patients that have already resolved SARS-CoV-2 infection. First of all, there was no difference in the frequency of early adaptive NK cells (NKG2C^+^CD57^-^) between HCMV^-^ REC patients in comparison to HCMV^-^ healthy donors (HC) that were sampled before the SARS-CoV-2 pandemic **(Figure 6H**, **left panel)**. Furthermore, the HCMV^-^ REC patients did not show any indication for expansion of mature adaptive NK cells (NKG2C^+^CD57^+^): none of the recovered patients did show an elevated frequency of mature adaptive NK cells (mean frequency 1.6%, ± 0.96%) compared to the HC cohort (mean frequency 1.3% ± 1.0%) **(figure 6H**, **right panel)**. This was compatible with HLA-DR expression of HCMV^-^ REC patients being low and comparable to that in HCMV^-^ HC patients **(figure 6D)**, illustrating that activation of early adaptive NK cells is restricted to the acute phase of infection. Similarly, the HCMV^+^ patients of the REC cohort did not have higher frequencies of NKG2C^+^CD57^+^ NK cells or increased levels of NKG2C compared to the HCMV^+^ donors in the HC cohort **(Supplementary Figure S10)**. Together, the data suggest that SARS-CoV-2 infection triggers activation and expansion of early adaptive NK cells but that the response is transitory and does not translate into the generation of a long-living population of mature adaptive NK cells.

## DISCUSSION

HLA-E is a central checkpoint for NK cell activation and its surface expression is majorly regulated by the differential availability of stabilizing peptides. Besides the endogenous peptides, typically encoded by HLA class I leader peptides, HLA-E is also efficiently upregulated in the course of HCMV infection by an immunodominant peptide processed from the UL40 viral protein. This is thought to represent an immuno-evasive measure of the virus, leading to inhibition of canonical NKG2A^+^ NK cells, which constitute the large majority of tissue-resident NK cells and which are also highly represented among circulating NK cells (*52*, *53*). On the other hand, HCMV infection triggers the generation of adaptive NK cells that are able to counteract this evasive mechanism by recognition of infected HLA-E-expressing cells via the stimulatory NKG2C receptor. The HLA-E-mediated balance between canonical and adaptive NK cells is thus key to the virus-specific immune response. The present study establishes SARS-CoV-2 as a further virus besides HCMV, that specifically increases HLA-E expression on infected cells by provision of an immunodominant peptide leading to inhibition of NKG2A^+^ canonical NK cells and stimulation of NKG2C^+^ adaptive NK cells. Moreover, analysis of COVID-19 patients in the acute phase of infection revealed that adaptive NK cells are developing and expanding independently of previous HCMV infection, a unique observation underlining the general relevance of adaptive NK cell responses in SARS-CoV-2 infection.

Our study demonstrates that productive infection with SARS-CoV-2 induces significantly increased cell surface expression levels of HLA-E in both primary human lung cells and lung epithelial cell lines. We could attribute this effect to a peptide encoded within the spike protein that increased HLA-E expression to a similar level than a peptide from the HLA-G leader sequence, representing one of the strongest known stabilizing peptides for HLA-E (*17*, *20*, *21*, *23*). Induction of HLA-E expression in TAP-deficient T2 cells was dose-dependent and strongest in T2 cells expressing HLA-E*01:03 compared to the other common allotype HLA-E*01:01, which is compatible with the generally stronger surface expression of HLA-E*01:03 (*17*). Furthermore, we could demonstrate that a single point mutation in the spike protein, changing the respective peptide sequence at position P5, completely abrogated the increase in HLA-E expression. This strongly suggests that the spike-mediated increase of HLA-E in human lung cells is predominantly due to efficient processing and presentation of the YLQPRTFLL peptide. By using a spike-GFP fusion protein, we could further demonstrate that the increase of HLA-E strictly correlates with the amount of spike protein and is not due to experimental stressors such as the transfection procedure or unspecific changes in cytokine levels.

The YLQPRTFLL site, whose *in silico* prediction was based on analysis of the original 2019 Wuhan strain sequence, was found to be present in all subsequently emerged WHO-defined SARS-CoV-2 variants of concern (VOCs) including the VOCs that successfully spread on a global scale (see GISAID database at gisaid.org). Although escape mutations were reported, they remained episodic and locally confined (*54*). Conservation of this peptide site might thus be of general structural and/or functional relevance for the spike protein, prohibiting fixation of immune evading mutations. Of note, the YLQPRTFLL peptide is also a known HLA-A*02-restricted T cell epitope and specific CD8^+^ T cells were identified in nearly 80% of HLA-A*02 patients infected with SARS-CoV-2 or vaccinated with a spike mRNA vaccine, but not in naïve individuals (*54–58*). More recently, HLA-E loaded with the YLQPRTFLL peptide was shown to elicit strong responses of HLA-E YLQPRTFLL-specific CD8+ T cell clones (*46*). The immunodominant nature of the peptide in the T cell context provides further compelling evidence that the peptide is properly and efficiently processed from the spike protein *in vivo*. Furthermore, the fact that this immunogenic site remains present in the major SARS-CoV-2 variants in spite of efficient recognition by NK as well as T cells argues for general constraints that prohibit any easy immune evasion mechanism of the virus.

Our observation that adaptive NK cells are activated during acute SARS-CoV-2 infection has several implications for NK cell-mediated immunity against this respiratory virus. Generally, ACE2-expressing lung cells represent a primary entry route for SARS-CoV-2 and our experiments illustrate how infected lung cells expressing high levels of ACE2 rapidly and strongly upregulate HLA-E. Notably, adaptive NK cells are present in the lung and expansions are found in the lung of healthy donors at comparable frequencies to peripheral blood. In particular, CD49a^+^NKG2C^+^ NK cells with a tissue-resident signature and a hyperresponsive functional state were found to be present in the lung (*59*). Thus, adaptive NK cells are already present on site, able to encounter SARS-CoV-2 at an early phase of infection in the lung and in the circulation and potentially contribute to virus-specific immunity by recognition of HLA-E on infected cells. According to the previously accepted paradigm, NKG2C^+^ adaptive NK cells are restricted to HCMV^+^ subjects, which in turn implied that only HCMV^+^ patients would be able to mount strong adaptive NK cell responses. The present study however suggests that this is not the case and that adaptive NK cell responses are mounted in acute SARS-CoV-2 infection irrespective of the HCMV status.

One of the most interesting aspects of this study is the observation that the proliferative response of adaptive NK cells during SARS-CoV-2 infection was largely restricted to early adaptive NK cells lacking CD57 expression. Although a minor subset among NKG2C^+^ NK cells in peripheral blood at steady-state, they majorly provoked the dynamic changes in the overall frequency of adaptive NK cells observed in a cohort of hospitalized COVID-19 patients. The fact that frequencies of early adaptive NK cells were back to baseline in HCMV^-^ patients that had recently recovered from COVID-19, makes it likely that the observed expansions of early adaptive NK cells in the acute phase of infection are transitory. The concomitant absence of increased frequencies of mature NKG2C^+^CD57^+^ adaptive NK cells in recovered COVID-19 patients makes it also unlikely that early adaptive NK cells converted into long-living mature adaptive NK cells. Our data thus suggest that SARS-CoV-2 infection including severe COVID-19 cases leads to a pronounced but transient increase of early NKG2C^+^CD57^-^ adaptive NK cells that is independent of the HCMV status. The reason for the failure to generate mature adaptive NK cells might be related to the fact that SARS-CoV-2 is typically cleared following the acute infection phase, whereas a latent phase is established following acute HCMV infection (*60*). The transition from early to mature adaptive NK cells seems to be a process that requires a latent phase of infection with continued presence of the virus, including sporadic events of reactivation, as is the case for HCMV. In this regard it will be interesting to look for the expansion of mature adaptive NK cells in cases where SARS-CoV-2 infection could not be rapidly cleared, as reported for patients with post-acute sequelae of COVID-19 (PASC), also known as Long-COVID (*61*, *62*). Indeed, increased frequencies of NKG2C^+^CD57^+^ mature adaptive NK cells were reported in a cohort of Long-COVID patients, even though the contribution of HCMV was not analyzed in this study (*63*).

What is the contribution of adaptive NK cells to the clinical course of COVID-19? On the one hand, adaptive NK cell frequencies are increased in severe acute COVID-19 disease as mentioned before (*42*, *64*, *65*). On the other hand, an increased risk of developing COVID-19 following SARS-CoV-2 infection was observed in a longitudinal study in subjects with low baseline frequencies of adaptive NKG2C^+^ NK cells (*66*). The same group in a separate study reported an association of high frequencies of canonical NKG2A^+^ and low frequencies of adaptive NKG2C^+^ NK cells with increased mortality in severe COVID-19 (*67*). A benefit of NKG2C^+^ NK cells in COVID-19 was also suggested by a genetic association study where hospitalized patients with severe symptoms of COVID-19 show an overrepresentation of the deleted allele of NKG2C (*KLRC2del*) and of the low expressing *HLA-E*01:01* allele compared to patients with mild symptoms (*68*). Indeed, in our study the YLQPRTFLL peptide was less efficiently presented by HLA-E*01:01, which is consistent with inferior recognition of SARS-CoV-2-infected cells by adaptive NK cells in patients lacking the high affinity allele *HLA-E*01:03*.

The present study shows that both, HCMV^-^ and HCMV^+^ subjects, are able to generate and expand early adaptive NK cells in response to SARS-CoV-2 infection. However, only HCMV^+^ subjects have preformed expansions of mature adaptive NK cells. A hypothesis emerging from the available data is that adaptive NK cells are generally beneficial to mount an early antiviral response and provide improved resistance against infection in the population. However, in the scenario of severe acute COVID-19, preformed large clonal expansions of mature adaptive NK cells have the potential to aggravate the disease by increased cytotoxicity as well as increased release of cytokines, thereby worsening systemic inflammation. Given the exquisite sensitivity of mature adaptive NK cells to IgG antibody-mediated activation, high IgG titers typically building up during acute severe infection could sustain their stimulatory response. It remains to be determined, if pre-existing expansions of mature adaptive NK cells in HCMV^+^ patients are associated with at greater risk to develop severe COVID-19.

In the ongoing SARS-CoV-2 vaccination programs that make use of RNA, DNA, or protein-based vaccines, the spike protein encoding the HLA-E binding peptide YLQPRTFLL is used as key immunogen (*69*). The present data implicate that HLA-E could be upregulated in antigen presenting cells (APCs) that internalize the vaccine, for example the spike-encoding RNA in case of the frequently used RNA vaccines. In this case, spike-peptide presenting APC, in particular dendritic cells, would be expected to accumulate at the injection site and in draining lymph node tissues (*70*). Increased HLA-E expression on APCs together with local and systemic spike-specific antibody responses elicited during the vaccination process constitute potent stimuli for adaptive NK cells. Although it is so far unknown whether HLA-E expression is indeed upregulated on APCs in response to SARS-CoV-2 vaccination, the role of early and mature adaptive NK cells, also in the context of repeated booster vaccinations, merits further investigation.

In summary, we demonstrate that SARS-CoV-2 induces HLA-E expression in infected cells and suggest that the spike-encoded peptide YLQPRTFLL is a main contributor to this effect. The upregulation of HLA-E differentially affects the NK cell-mediated recognition of virus-infected cells. On the one hand, adaptive NK cells are strongly stimulated via the HLA-E-NKG2C axis. Due to concomitant downregulation of classical HLA class I in the infected cells, additional stimulatory input is provided by the lack of inhibition of the cognate inhibitory KIR, which is generally co-expressed on adaptive NK cells together with NKG2C. On the other hand, canonical NKG2A^+^ NK cells, which mostly do not express a cognate KIR, are strongly inhibited by the complex of HLA-E and spike peptide. Analysis of patients with severe COVID-19 revealed that during the acute phase of infection, the functional balance between canonical and adaptive NK cells is tipped due to the generation and activation of NKG2C^+^CD57^-^ adaptive NK cells. Unexpectedly, expansions of these early adaptive NK cells occurred not only in HCMV^+^ patients already having mature adaptive NK cells but also in HCMV^-^ patients. The selective expansion of early but not mature adaptive NK cells is a unique observation so far not made in other virus infections and might constitute an important novel and clinically relevant aspect of the immune response against SARS-CoV-2.

