## Supplementary figures 4-10 for "SARS-CoV-2 modulates NK cell responses via induction of HLA-E and triggers expansion of adaptive NK cells during acute infection"

### Slide 1
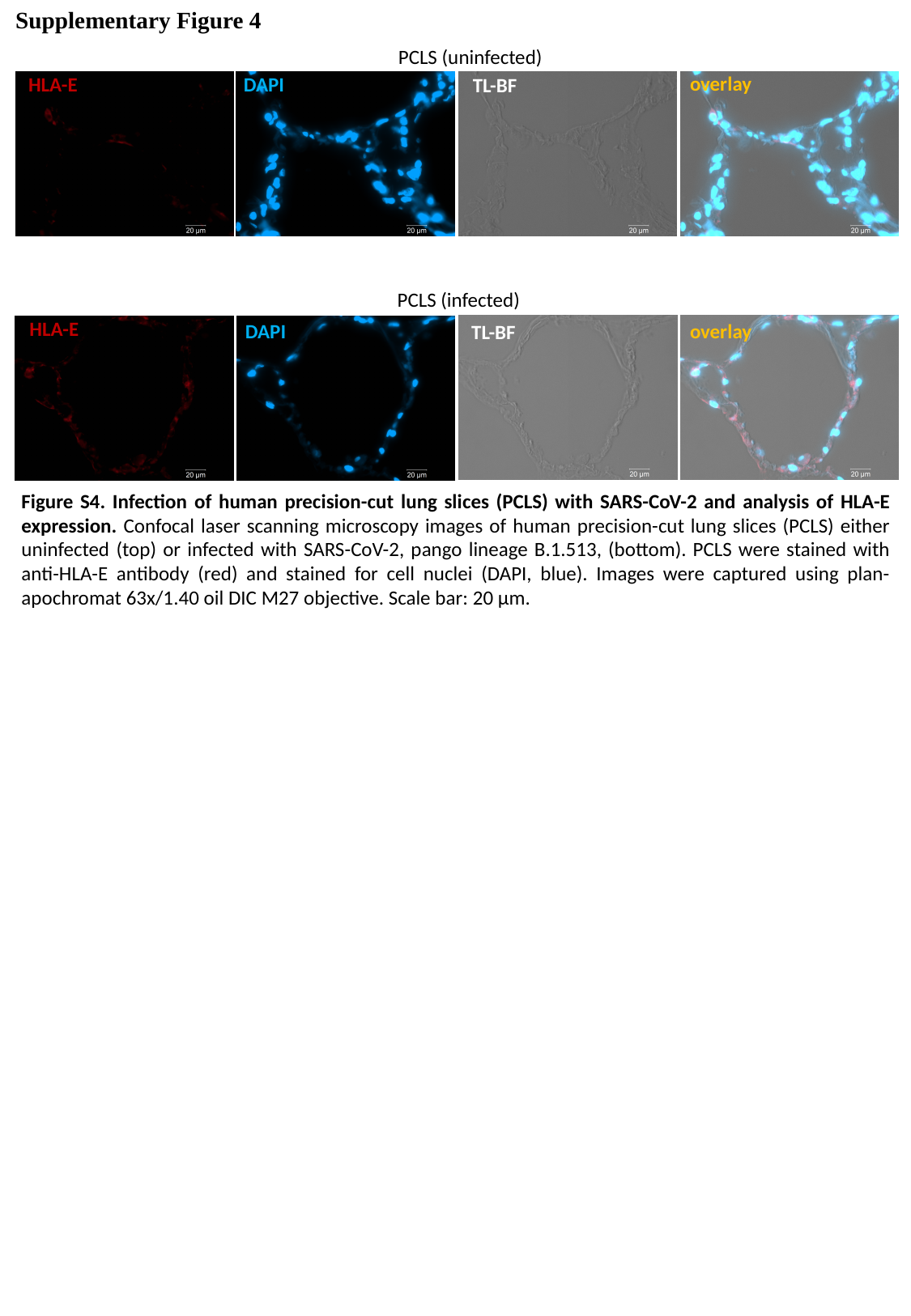

Supplementary Figure 4
PCLS (uninfected)
overlay
DAPI
HLA-E
TL-BF
PCLS (infected)
HLA-E
overlay
DAPI
TL-BF
Figure S4. Infection of human precision-cut lung slices (PCLS) with SARS-CoV-2 and analysis of HLA-E expression. Confocal laser scanning microscopy images of human precision-cut lung slices (PCLS) either uninfected (top) or infected with SARS-CoV-2, pango lineage B.1.513, (bottom). PCLS were stained with anti-HLA-E antibody (red) and stained for cell nuclei (DAPI, blue). Images were captured using plan-apochromat 63x/1.40 oil DIC M27 objective. Scale bar: 20 µm.

### Slide 2
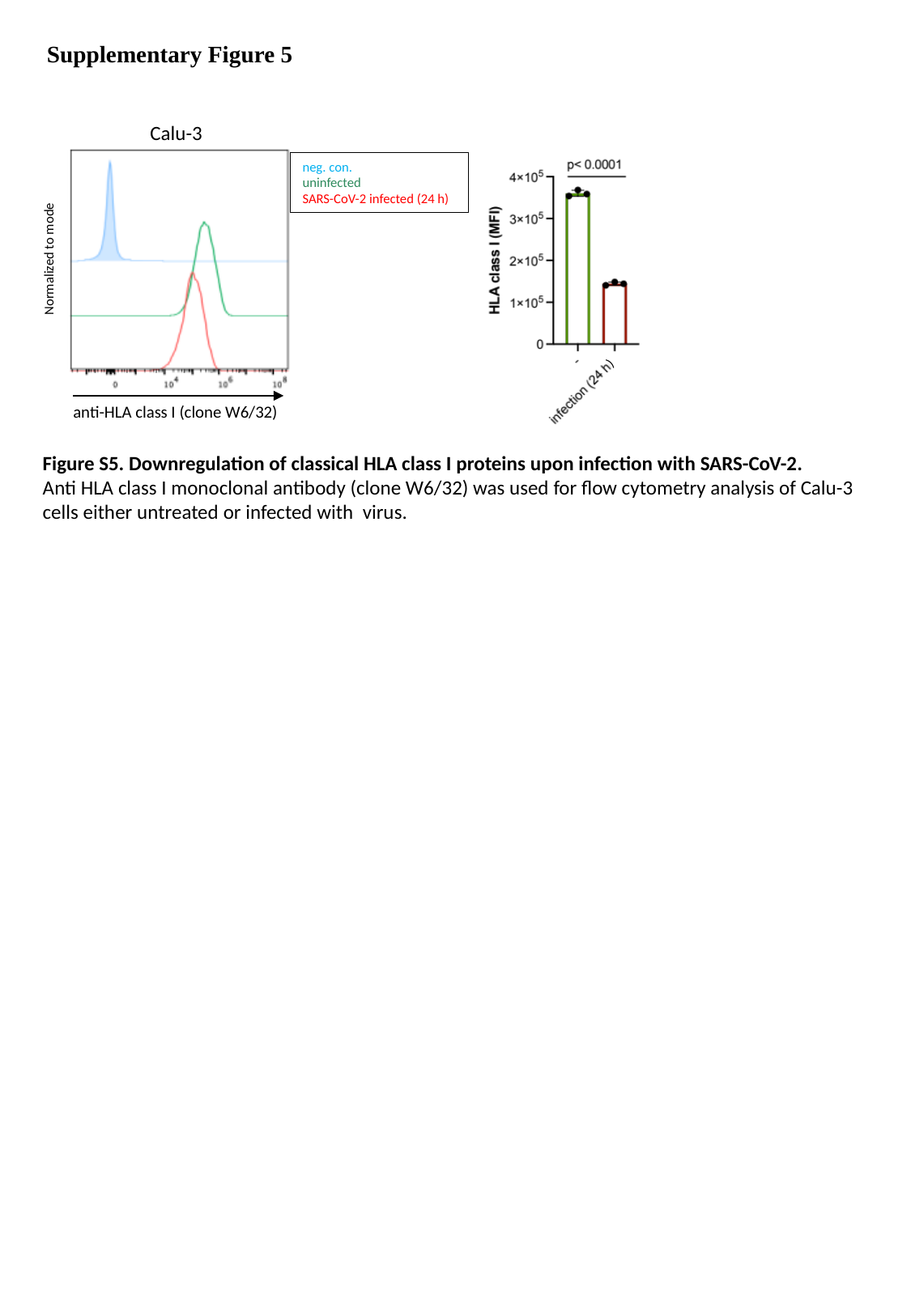

Supplementary Figure 5
Calu-3
neg. con.
uninfected
SARS-CoV-2 infected (24 h)
Normalized to mode
)
anti-HLA class I (clone W6/32)
Figure S5. Downregulation of classical HLA class I proteins upon infection with SARS-CoV-2.
Anti HLA class I monoclonal antibody (clone W6/32) was used for flow cytometry analysis of Calu-3 cells either untreated or infected with virus.

### Slide 3
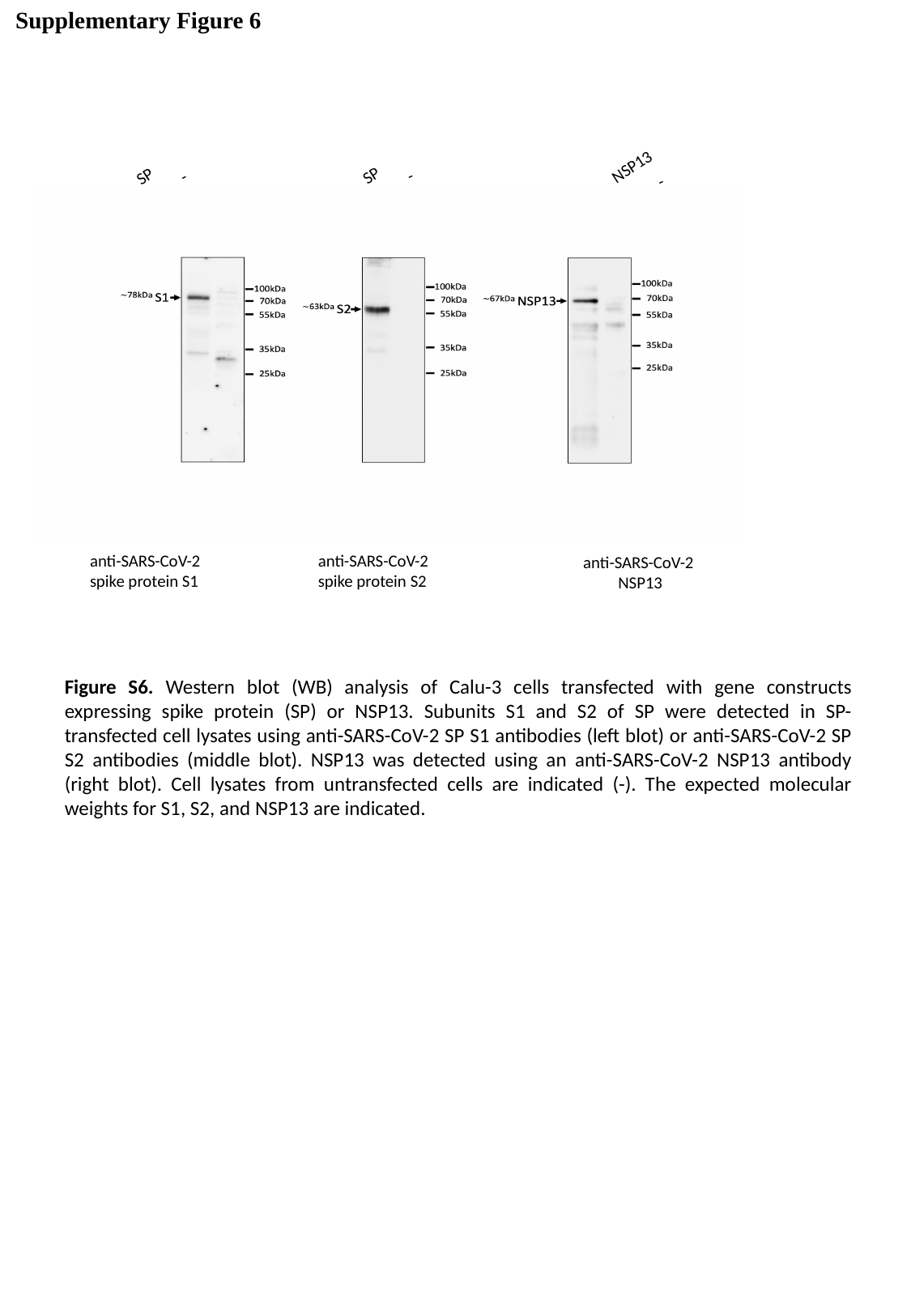

Supplementary Figure 6
NSP13
SP
-
SP
-
-
anti-SARS-CoV-2
spike protein S2
anti-SARS-CoV-2
spike protein S1
anti-SARS-CoV-2
NSP13
Figure S6. Western blot (WB) analysis of Calu-3 cells transfected with gene constructs expressing spike protein (SP) or NSP13. Subunits S1 and S2 of SP were detected in SP-transfected cell lysates using anti-SARS-CoV-2 SP S1 antibodies (left blot) or anti-SARS-CoV-2 SP S2 antibodies (middle blot). NSP13 was detected using an anti-SARS-CoV-2 NSP13 antibody (right blot). Cell lysates from untransfected cells are indicated (-). The expected molecular weights for S1, S2, and NSP13 are indicated.

### Slide 4
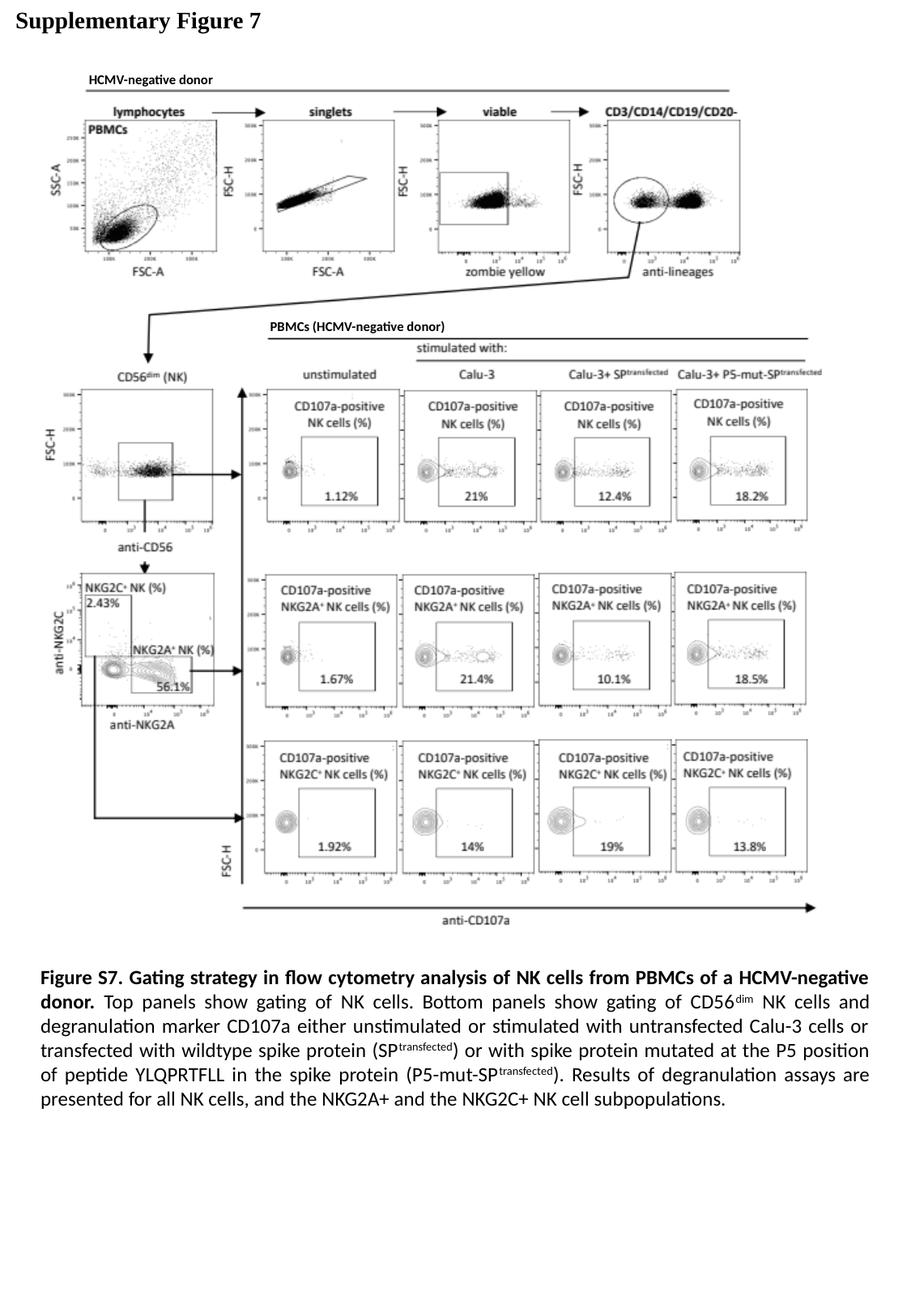

Supplementary Figure 7
HCMV-negative donor
PBMCs (HCMV-negative donor)
Figure S7. Gating strategy in flow cytometry analysis of NK cells from PBMCs of a HCMV-negative donor. Top panels show gating of NK cells. Bottom panels show gating of CD56^dim^ NK cells and degranulation marker CD107a either unstimulated or stimulated with untransfected Calu-3 cells or transfected with wildtype spike protein (SP^transfected^) or with spike protein mutated at the P5 position of peptide YLQPRTFLL in the spike protein (P5-mut-SP^transfected^). Results of degranulation assays are presented for all NK cells, and the NKG2A+ and the NKG2C+ NK cell subpopulations.

### Slide 5
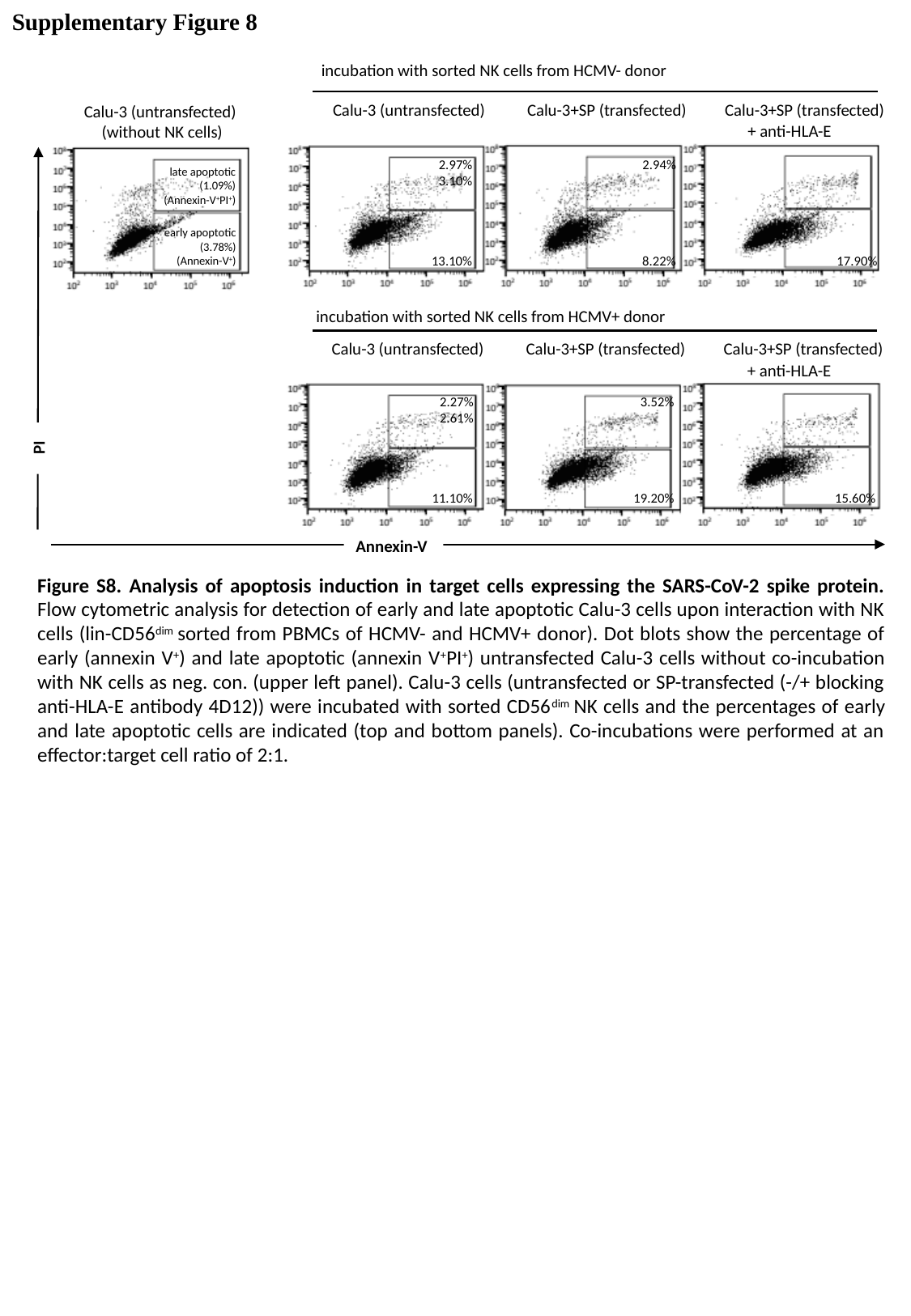

Supplementary Figure 8
incubation with sorted NK cells from HCMV- donor
Calu-3 (untransfected) Calu-3+SP (transfected) Calu-3+SP (transfected)
Calu-3 (untransfected)
(without NK cells)
+ anti-HLA-E
2.97% 2.94% 3.10%
late apoptotic
(1.09%)
(Annexin-V^+^PI^+^)
early apoptotic
 (3.78%)
(Annexin-V^+^)
13.10% 8.22% 17.90%
incubation with sorted NK cells from HCMV+ donor
Calu-3 (untransfected) Calu-3+SP (transfected) Calu-3+SP (transfected)
+ anti-HLA-E
2.27% 3.52% 2.61%
 PI
11.10% 19.20% 15.60%
Annexin-V
Figure S8. Analysis of apoptosis induction in target cells expressing the SARS-CoV-2 spike protein. Flow cytometric analysis for detection of early and late apoptotic Calu-3 cells upon interaction with NK cells (lin-CD56^dim^ sorted from PBMCs of HCMV- and HCMV+ donor). Dot blots show the percentage of early (annexin V^+^) and late apoptotic (annexin V^+^PI^+^) untransfected Calu-3 cells without co-incubation with NK cells as neg. con. (upper left panel). Calu-3 cells (untransfected or SP-transfected (-/+ blocking anti-HLA-E antibody 4D12)) were incubated with sorted CD56^dim^ NK cells and the percentages of early and late apoptotic cells are indicated (top and bottom panels). Co-incubations were performed at an effector:target cell ratio of 2:1.

### Slide 6
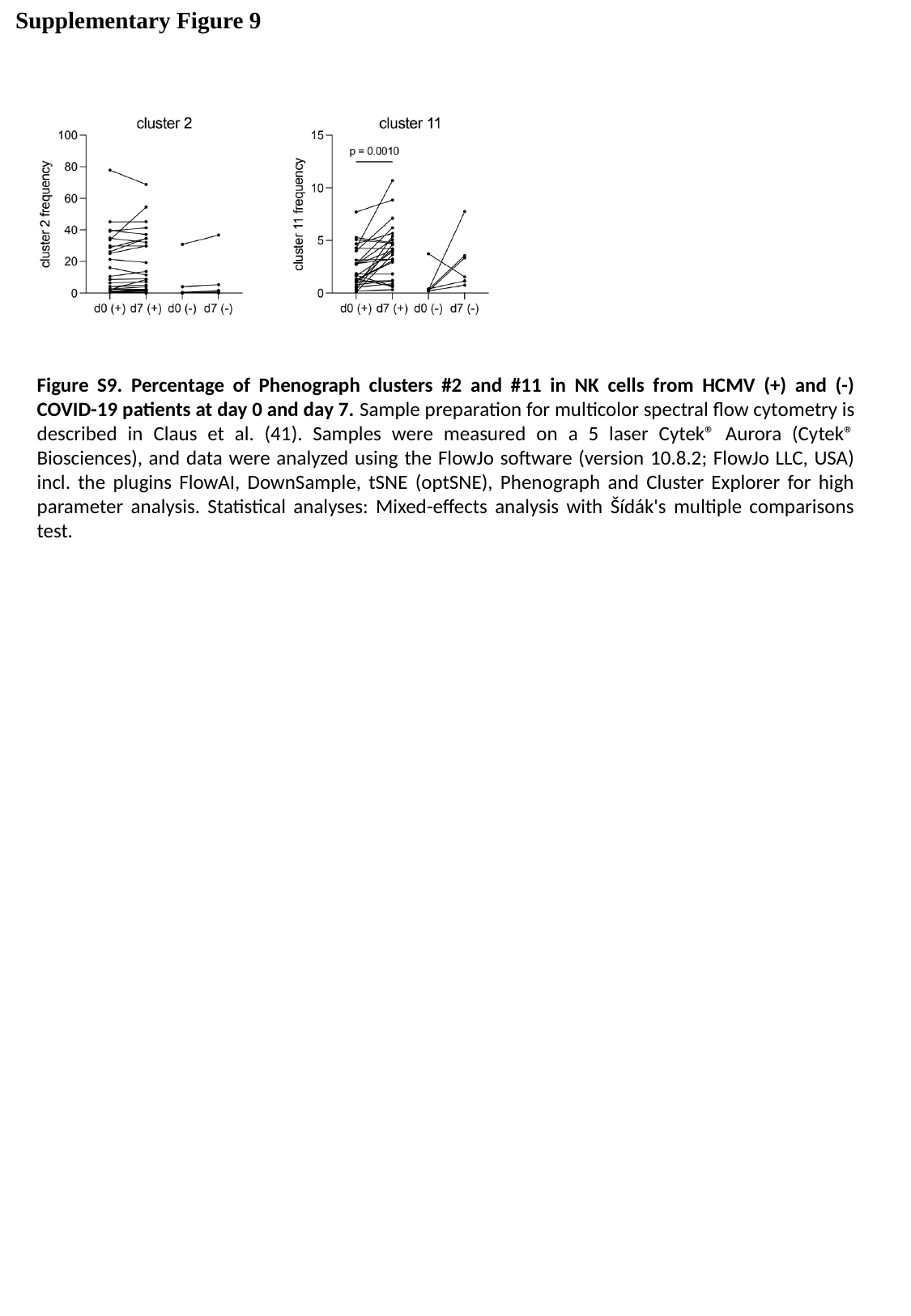

Supplementary Figure 9
Figure S9. Percentage of Phenograph clusters #2 and #11 in NK cells from HCMV (+) and (-) COVID-19 patients at day 0 and day 7. Sample preparation for multicolor spectral flow cytometry is described in Claus et al. (41). Samples were measured on a 5 laser Cytek® Aurora (Cytek® Biosciences), and data were analyzed using the FlowJo software (version 10.8.2; FlowJo LLC, USA) incl. the plugins FlowAI, DownSample, tSNE (optSNE), Phenograph and Cluster Explorer for high parameter analysis. Statistical analyses: Mixed-effects analysis with Šídák's multiple comparisons test.

### Slide 7
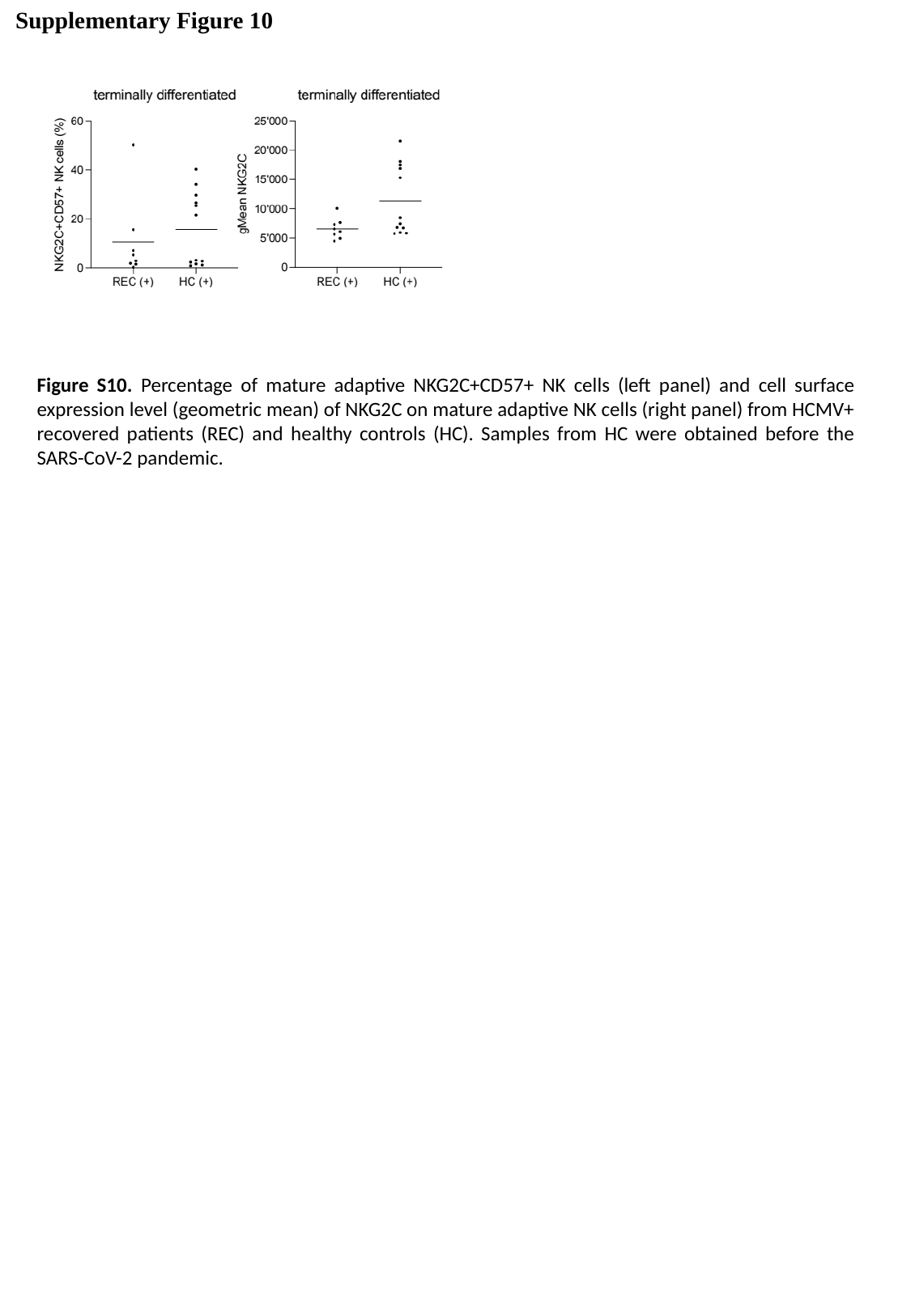

Supplementary Figure 10
Figure S10. Percentage of mature adaptive NKG2C+CD57+ NK cells (left panel) and cell surface expression level (geometric mean) of NKG2C on mature adaptive NK cells (right panel) from HCMV+ recovered patients (REC) and healthy controls (HC). Samples from HC were obtained before the SARS-CoV-2 pandemic.
